# Temporal transcriptional rhythms govern coral-symbiont function and microbiome dynamics

**DOI:** 10.1101/2025.08.06.668741

**Authors:** Bradley Allen Weiler, Nicholas Kron, Anthony Mario Bonacolta, Mark J.A. Vermeij, Andrew Charles Baker, Javier del Campo

## Abstract

Diel rhythms align physiological processes with light/dark cycles, driving predictable oscillations in gene expression and protein activity through tightly controlled transcriptional-translational feedback loops. Transcriptomic analyses in the stony coral *Pseudodiploria strigosa* revealed tightly regulated transcriptional control: dawn triggers a molecular reset marked by RNA metabolism and protein turnover; midday emphasizes anabolic and phosphate-regulated pathways; dusk reflects transitional lipid and amino acid metabolism; and midnight reveals stress-responses, mRNA catabolism, and mitochondrial organization. The photosymbiont *Breviolum* sp. exhibits subtler but distinct diel regulation, with photoprotective activation at dawn, metabolite transport and nitrogen cycling through midday and dusk, and cell cycle and ion homeostasis at night. Microbial communities show time-dependent restructuring of co-occurrence networks. Carbohydrate-catabolizing taxa associated with Symbiodiniaceae during daylight, while nighttime assemblages feature methylotrophs likely facilitating dimethylsulfoniopropionate (DMSP) metabolism. Together, these findings present the first system-level molecular framework of diel regulation across the coral-photosymbiont-microbe holobiont, revealing strong time-specific transcriptional control as a key driver of coordinated function and homeostasis.

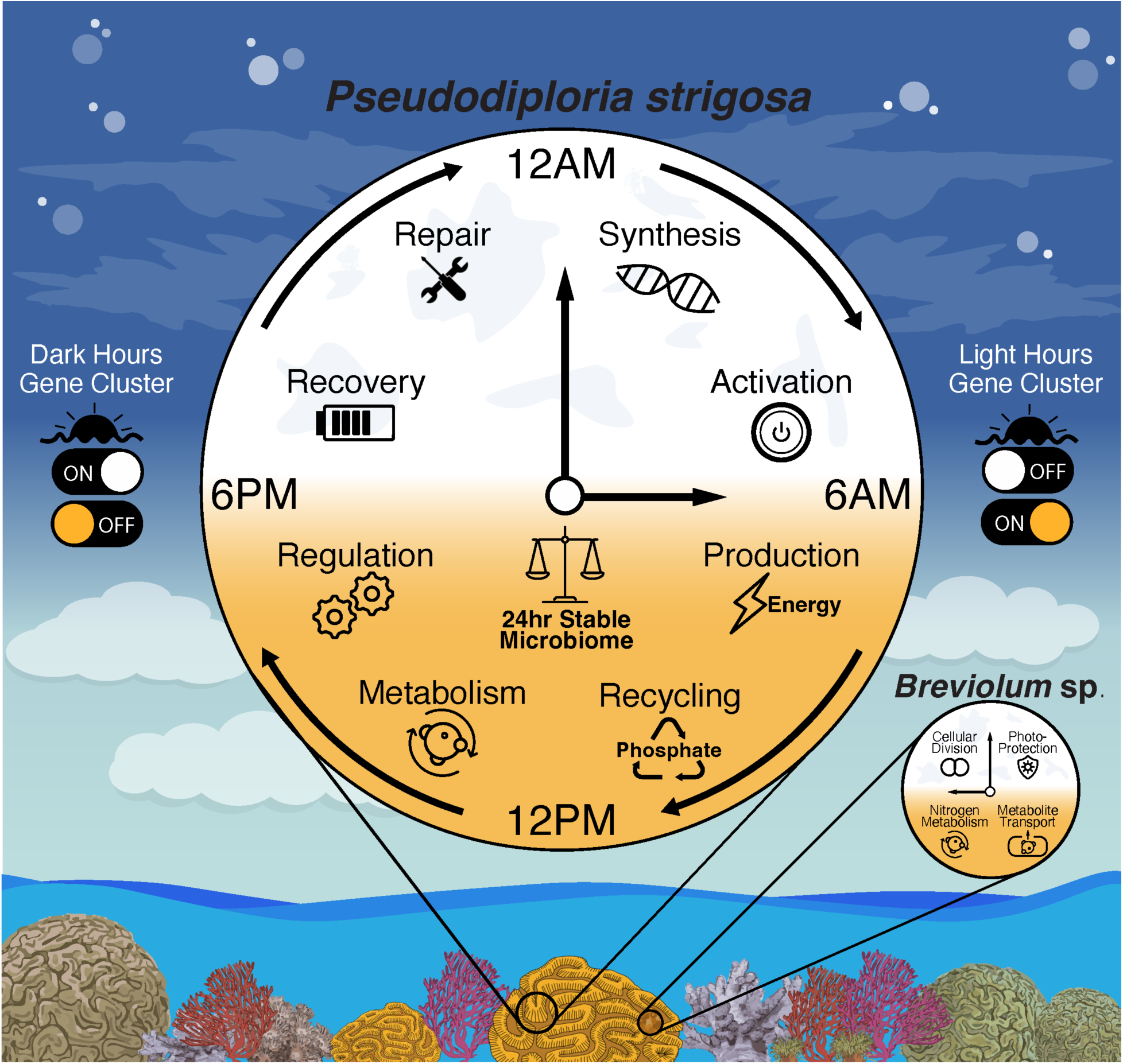

## Introduction

Natural light-dark cycles shape core cellular mechanisms that govern homeostasis and key metabolic functions ^1–3^. Diel cycles entrain organismal circadian rhythms, orchestrating predictable oscillations in gene expression and protein activity through transcriptional-translational feedback loops ^4,5^. Central to these rhythms are three interconnected modules: sensory elements (i.e., light-sensing photoreceptors) that interpret environmental cues; autoregulatory transcriptional feedback loops that generate rhythmic signals; and clock-controlled genes (CCGs) that regulate downstream cellular processes. While circadian rhythms are often studied in the context of mammalian sleep, recent work has uncovered rhythmic activity cycles, functionally analogous to rest and activity cycles in invertebrates lacking centralized nervous systems, including placozoans ^6^, hydra ^7^, sponges ^8^, and jellyfish ^9^. These findings suggest that temporal regulation of physiological states may be an ancient feature of multicellular life and may also extend to host-microbe interactions ^10^.

Scleractinian coral physiology is intrinsically linked to their microbiome, a diverse community of bacteria, archaea, fungi, protists, algae, and viruses that collectively maintain holobiont homeostasis ^11–14^. Symbiotic algae in the Family Symbiodiniaceae ^15,16^ play a crucial role in coral metabolism by increasing oxygen levels to ∼250% of air saturation, creating hyperoxic conditions ^17^, and translocating up to 80% of the coral’s daily carbon requirements as photosynthates during diel light cycles ^18^. Microbial associates also play key roles in biogeochemical cycles, including the catabolism of complex carbohydrates to support the holobiont energy demands ^19^. While the influence of diel cycles on coral-associated microbial composition remains poorly understood, observed temporal variation in microbial communities has generally been limited ^20–22^. Most studies lacked replicated, high-resolution sampling across multiple days, limiting the ability to detect rhythmic shifts in microbiome composition across light and dark cycles. Caughman et al. (2021) observed subtle diel shifts in Proteobacteria associated with *Porites* spp., including enriched frequency of *Aeromonas* and *Vibrio* during the day and *Porticoccus* at night, potentially linked to hydrocarbon cycling of volatile organic compounds produced by Symbiodiniaceae ^21,23^. Seiblitz et al. (2025) recovered two amplicon sequence variants (ASVs) in two stony corals with 12- and 24-hour periodicity (ASV477: *Woesia* sp. and ASV148: *Pseudoaltermonas* sp., respectively). Silveira et al. (2017) even observed no clear rhythmic trends, despite sampling stony coral mucus every six hours for two days, implying that the absence of strong rhythmic signal may reflect host specificity or limitations in study design more than true biological stability. As all holobiont members encode circadian machinery, resolving temporal shifts in microbial communities remains essential for understanding whether diel rhythmicity arises independently or through complex host-microbe interactions.

Diel regulation of gene expression has been documented using primarily lab-based experiments on corals including *Favia* ^24^, *Euphyllia* ^25^, and multiple *Acropora* species ^26–33^. These studies report oscillations in core circadian regulators: *clock*, cryptochromes 1 and 2 (*cry1*/*2*), *cycle* (BMAL1 ortholog), and in some cases period (*per1* and *per2*) and timeless (*tim*) ^34,35^. However, most prior studies focused on canonical clock genes and overlooked the broad suite of CCGs that facilitate time-specific physiological processes. A more integrated analysis across holobiont components is necessary to determine whether CCGs downstream of the core oscillators are rhythmically expressed in a coordinated manner. To address this, we used high-resolution time-series sampling over three natural diel cycles in the Caribbean coral *Pseudodiploria strigosa*, profiling gene expression in the host and its endosymbionts in the genus *Breviolum* (Family Symbiodiniaceae), alongside molecular surveys of the microbiome. We identified a suite of rhythmically expressed CCGs in the host (∼30% of filtered transcripts), including genes oscillating at six-hour intervals, indicating tightly regulated transcriptional control. Similarly, *Breviolum* exhibited rhythmic gene expression through time, although more muted than the host, while specific microbial taxa showed subtle but consistent temporal enrichment. These results indicate that invertebrates without centralized nervous systems contain deeply conserved cellular machinery calibrated to circadian rhythms, highlighting circadian coordination across domains as an ancient feature of multicellular life.

## STAR Methods

### KEY RESOURCES TABLE

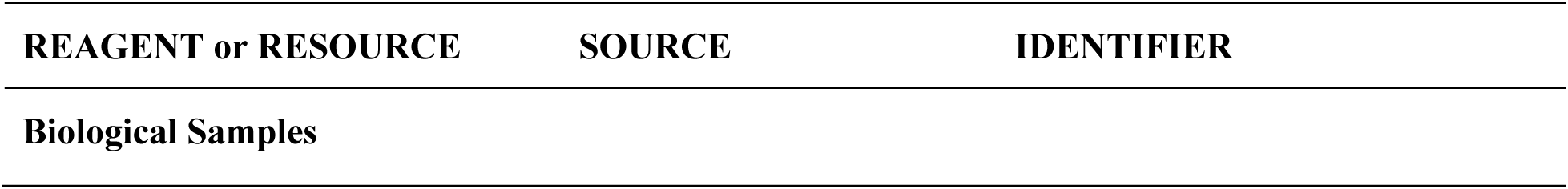
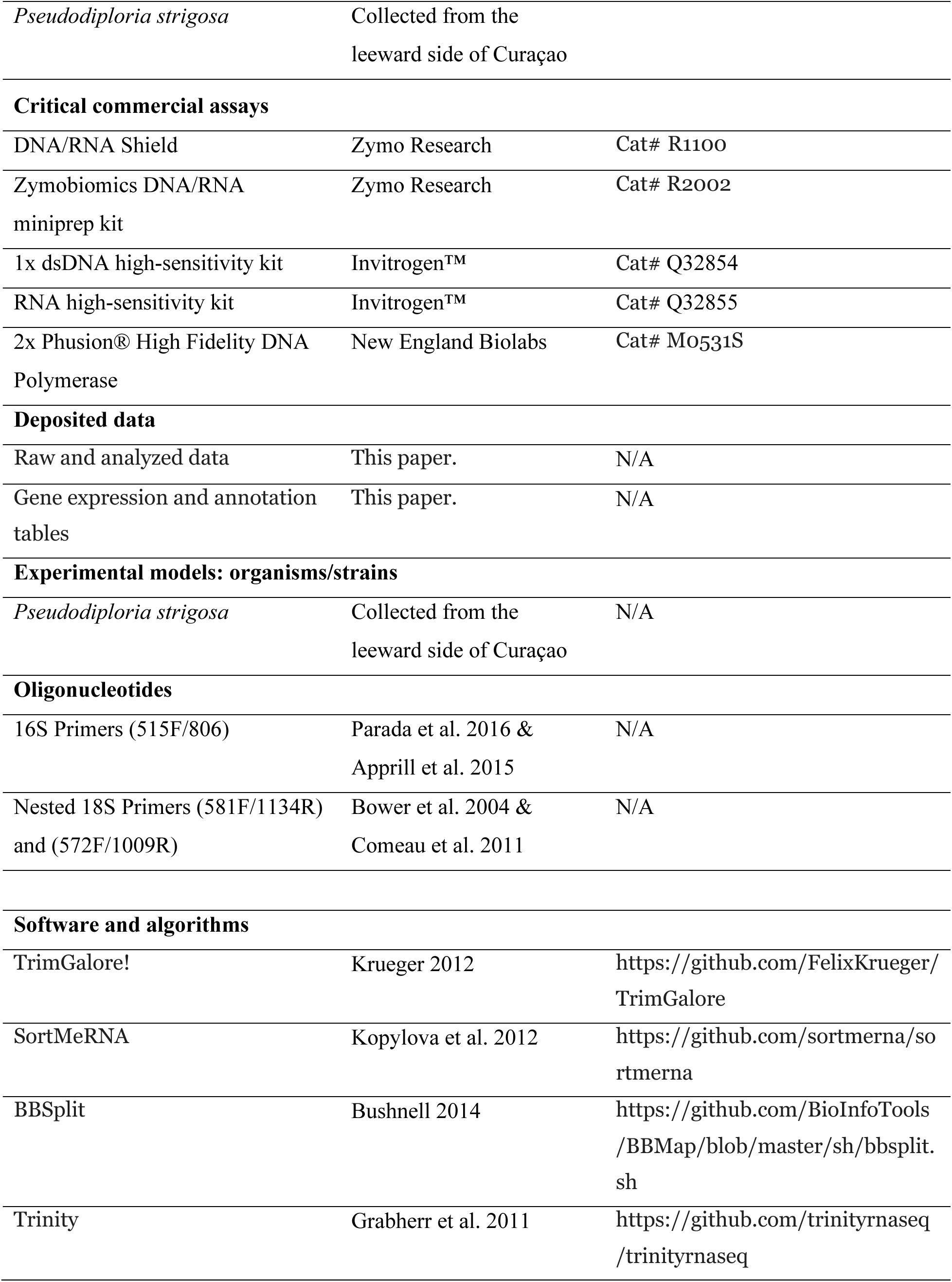
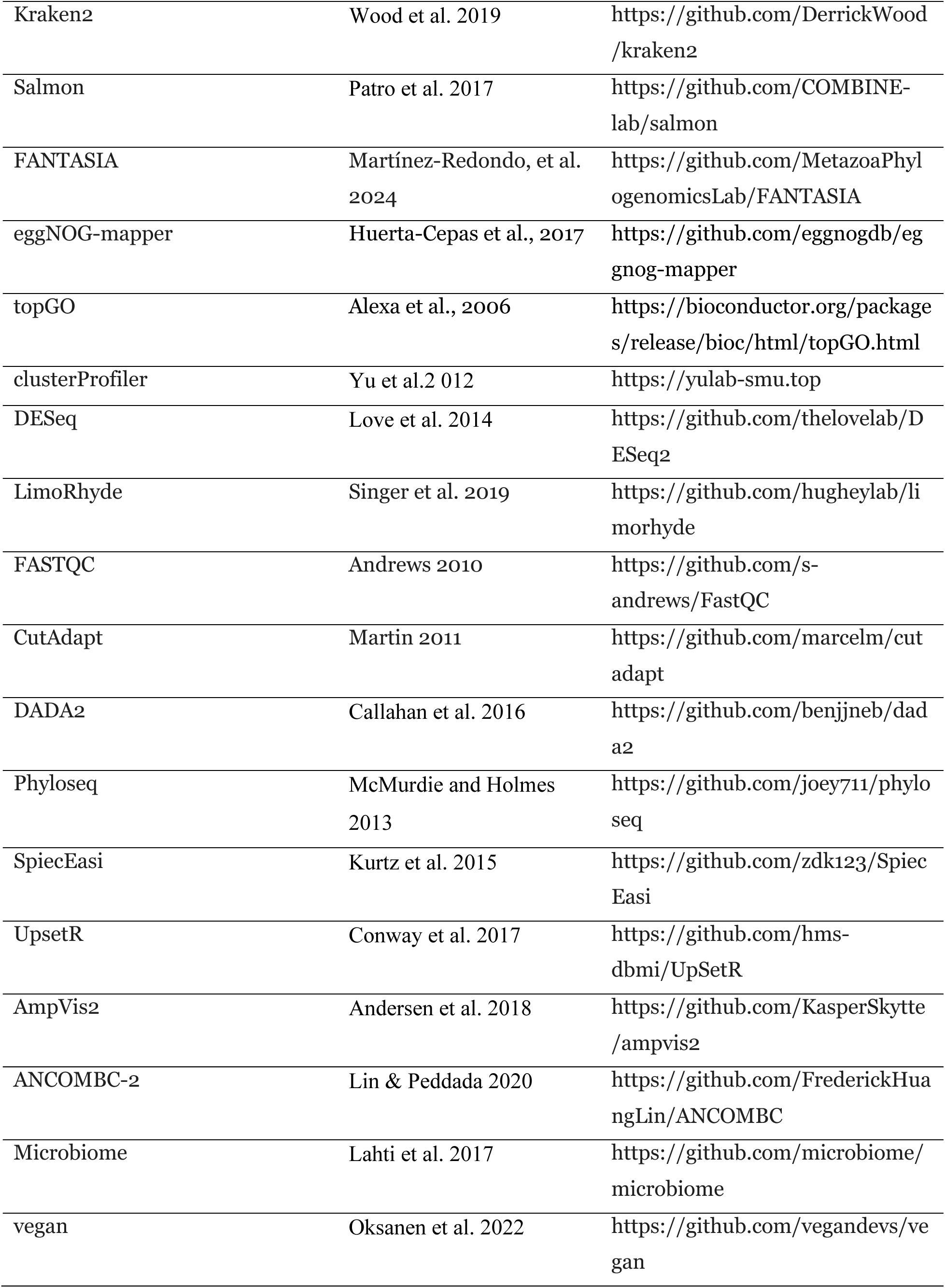
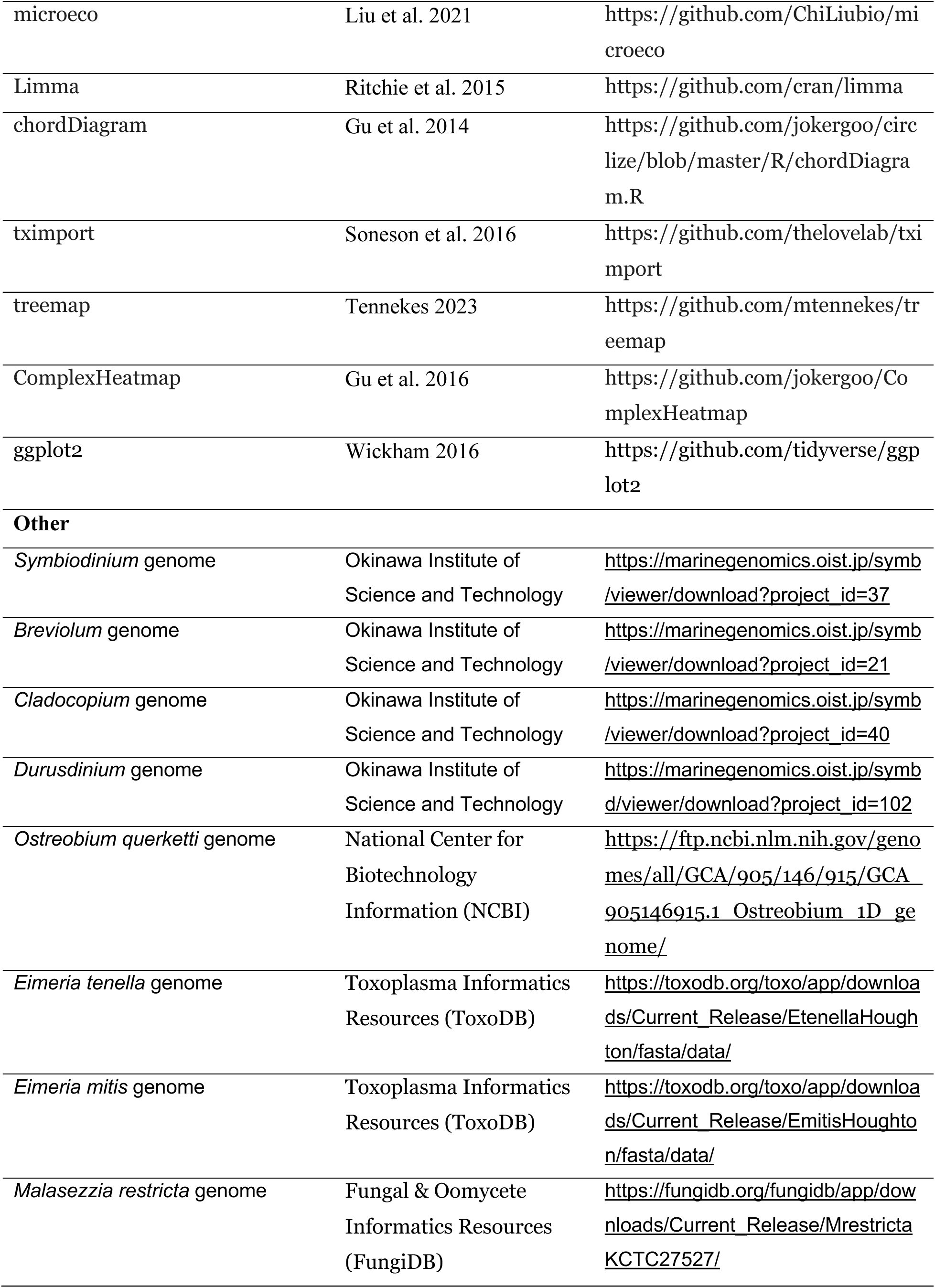
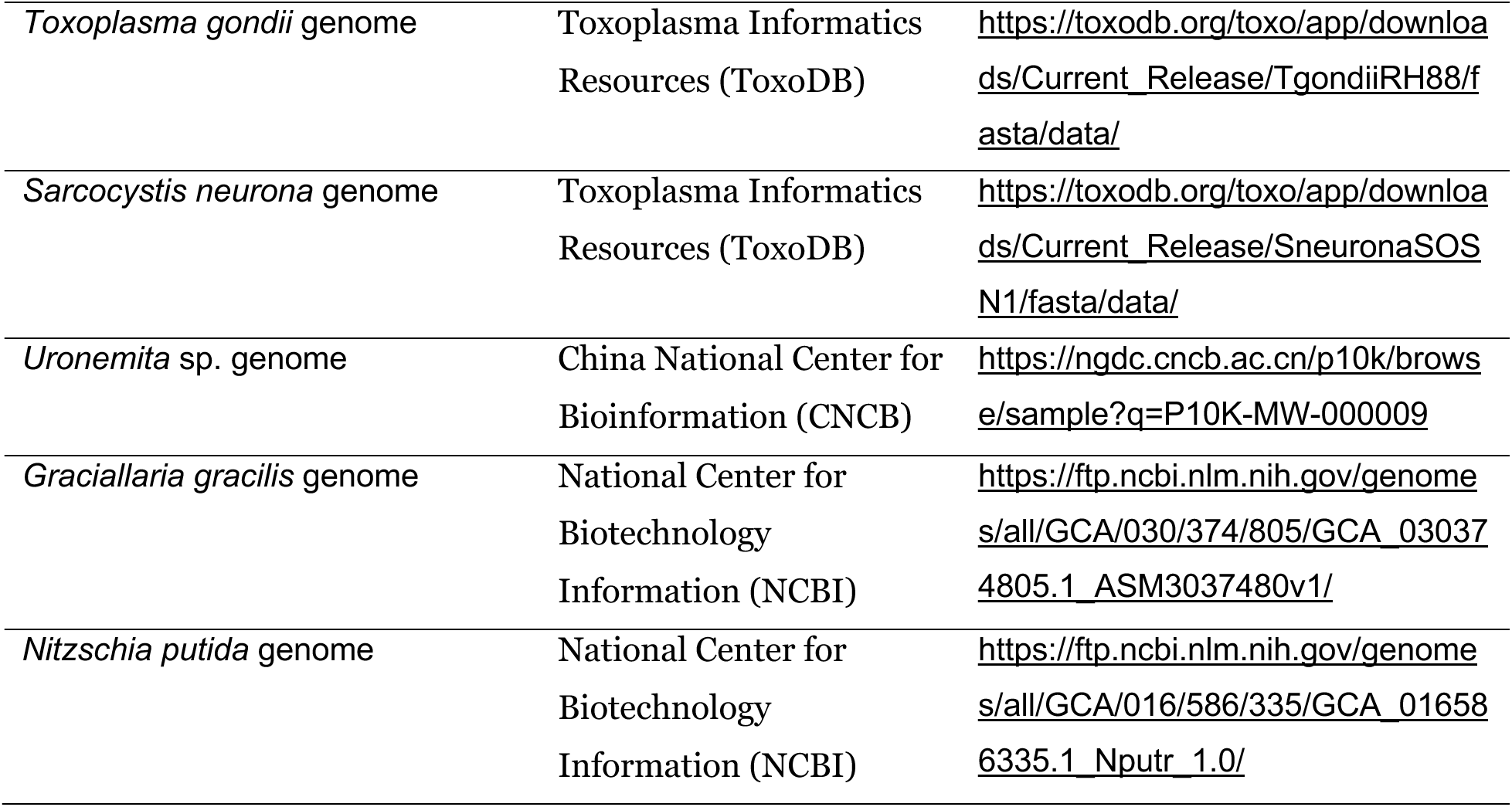

### RESOURCE AVAILABILITY

#### Lead contact

Further information and requests for resources and reagents should be directed to and will be fulfilled by the Lead Contact, Javier del Campo.

#### Materials availability

This study did not generate new, unique reagents.

#### Data and code availability

The raw reads for the project have been deposited on NCBI (SRA: XXXXXXXX). Code used for analysis, ASV count tables, taxonomy tables, and nucleotide sequences can be found on GitHub at: https://github.com/delCampoLab/diel_pstr.

### EXPERIMENTAL MODEL AND STUDY PARTICIPANT DETAILS

#### Coral diel sampling

Sampling was conducted in shallow (depth: ∼5m) water using SCUBA on the leeward side of Curaçao (12.120824, −68.969706; Supplementary Figure 1A&B) at six-hour intervals (n=4/day) starting at 12 AM (00:00 or midnight) on the 15^th^ of October and ending at 6 PM on the 17^th^ of October 2022 for a total of 12 sampling time-points. The light periods of the day during October closely followed a 12:12 light to dark period, with sunrise at 6:25 AM and sunset 6:18 PM. Sampling occurred within minutes of each timepoint described to limit sampling variability. Coral tissue biopsies (∼1cm^3^) were collected in triplicate from the same colony of scleractinian coral *Pseudodiploria strigosa* (Supplementary Figure 1C) using standard wire cutters at each time-point (n=36), stored in individually sealed bags (Uline, Florida, USA) and processed at the CARMABI marine research station (Collection Permit # 2022/021467). A reference water sample was also collected at each time point using a bleach-sterilized collapsible 1.5L Nalgene water bottle. Coral samples were stored in 1.2ml cryovials using flame-sterilized forceps and submerged in DNA/RNA Shield (Zymo Research, Irvine, CA, USA) for total nucleic acid preservation. One liter of the reference water samples was filtered over a 0.22 μm Isopore™ membrane filter (MilleporeSigma), and filters were submerged in cryovials with DNA/RNA shield. Samples were transported to the University of Miami Rosenstiel School of Marine, Earth, and Atmospheric Sciences for DNA/RNA co-extraction and long-term storage (CITES Permit #20US835702/9).

### METHODS DETAILS

#### Nucleic acid extraction and sequencing

All coral samples were co-extracted using the Zymobiomics DNA/RNA miniprep kit (Zymo Research, Irvine, CA, USA) with a slight modification to the manufacturer’s protocol, increasing the duration of bead-beating from 5 minutes to 40 minutes. Water samples were extracted for DNA using the same kit (not extracting for RNA); modifying the manufacturer’s protocol with a 10-minute bead beating step. All samples were quantified for DNA and RNA concentration using the Qubit™ 4.0 fluorometer (Invitrogen™, Carlsbad, CA, USA) 1x dsDNA high sensitivity kit (catalog # Q32854) and RNA high sensitivity kit (catalog #Q32855), respectively. Samples for 16S rRNA gene metabarcoding were diluted according to the sequencing facilities guidelines (Novogene Co. Ltd., Beijing, China), amplifying the V4 hypervariable region of the 16S rRNA gene with previously published primers 515F (5’-GTGYCAGCMGCCGCGGTAA ^36^) and 806R (5’-GGACTACNVGGGTWTCTAAT ^37^) using the Illumina NovaSeq paired-end (PE) 250bp.

18S rRNA gene metabarcoding samples used a nested PCR to exclude metazoan hosts prior to amplicon sequencing ^38^. In short, samples were amplified using non-metazoan primers EUK581-F (5’-GTGCCAGCAGCCGCG ^39^) and EUK1134-R (5’-TTTAAGTTTCAGCCTTGCG ^39^) targeting the V4 hypervariable region using 2x Phusion® High Fidelity DNA Polymerase (New England Biolabs, Ipswich, MA, USA) with a thermal cycler programmed at: 98°C for 30s, then 35 cycles of 98°C for 10s, 51.1°C for 30s, 72°C for 60s, with a final extension of 72°C for 5 minutes. Next, PCR products are validated on a 1% gel using electrophoresis for 60 minutes at 110 volts. Finally, the PCR products are sent to Novogene (Beijing, China) to be sequenced using the Illumina NovaSeq PE 250bp using previously published primers 572F (5’-CYGCGGTAATTCCAGCTC ^40^) and 1009R (5’-AYGGTATCTRATCRTCTTYG ^40^) amplifying the 18S rRNA gene V4 hypervariable region.

All samples were quantified for RNA concentration using the Qubit™ 4.0 fluorometer (Invitrogen™, Carlsbad, CA, USA) RNA high-sensitivity kit (catalog #Q32855), respectively. RNA sequencing was conducted using a Poly-A capture/enrichment methodology by Novogene (Beijing, China) on the Illumina NovaSeq PE 150bp. The methodology follows a standard workflow of fragmentation, reverse transcription, cDNA synthesis, end-repair and A-tailing, adapter ligation, size selection, PCR amplification, purification, and finally quantification using Qubit™ (Invitrogen™, Carlsbad, CA, USA) and RT-PCR and the size distribution validated with Agilent 2100 Bioanalyzer (Santa Clara, CA, USA).

### QUANTIFICATION AND STATISTICAL ANALYSIS

#### Bioinformatics and data analysis

For metabarcoding data from 16S and 18S rRNA genes, the raw reads were trimmed of adapter sequences by CutAdapt version 4.9 ^41^ and then quality checked using FastQC version 0.11.5 ^42^. Amplicon sequence variants were inferred through R package DADA2 version 1.26.0 ^43^, and taxonomy was assigned using DADA2 trained reference databases SILVA (non-redundant 99%) version 138.1 ^44^ for 16S and PR2 version 5.0.0 ^45^ for 18S. Phyloseq objects were compiled using the R package “Phyloseq” version 1.42.0 ^46^. Alpha/beta diversity and core communities were explored using the R package “Microbiome” ^47^ version 1.20.0. Community profiles and abundance plots were produced using R packages “ggplot2” ^48^ and “vegan” ^49^, while the differential abundance of ASVs between variables was conducted using “ANCOMBC” version 2.0.2 ^50^. Heatmaps for community composition were produced using “AmpVis2” version 2.8.6 ^51^, while the core community was plotted using “UpsetR” version 1.4.0 ^52^ for upset plots and “ggvenn” version 0.1.10 for Venn diagrams. Co-occurrence plots were generated using R package “microeco” version 1.1.0 ^53^, with correlation parameters set to SparCC, a filter threshold of 0.0001, and network building method using “SpiecEasi” version 1.1.3 ^54^.

RNA sequencing reads were first quality checked using FastQC version 0.11.5 ^42^; then adapters were trimmed using CutAdapt ^41^ wrapper script TrimGalore! ^55^, with a final quality check post-trim. Ribosomal RNA was partitioned from total RNA using SortMeRNA version 4.3.6 ^56^ and the default fasta reference database version 4.3. The remaining RNA was mapped to an index of genomes (*Ostreobium quekettii, Symbiodinium* sp. ^57^*, Breviolum* sp. ^58^*, Cladocopium sp.* ^57^*, Durusdinium* sp. ^59^*, Eimeria tenella, Emitis houghton, Sarcocystis neurona, Toxoplasma gondii, Malassezia restricta, Nitzschia putrida, Gracilaria gracilis,* and *Uronemita* sp.) to isolate non-coral RNA reads using BBsplit ^60^. Unmapped reads were concatenated for *de novo* transcriptome assembly of the various coral hosts using Trinity ^61^, and then decontaminated using the Standard PlusPF reference database in Kraken2 ^62^. Individual sample reads were aligned to the corresponding assembled transcriptome using Salmon’s pseudoalignment and quantification ^63^ for quantitative gene transcript expression analysis. STAR was used to align the *Breviolum* sp. binned reads from bbsplit to the *Breviolum* reference genome to produce quantitative files for exploring symbiont transcript expression. Gene transcripts were first predicted into longest open reading frames (ORFs) using transDecoder (https://github.com/TransDecoder/TransDecoder) and annotated using EggNOG-mapper ^64^ to assign genes to predicted proteins and orthologous groups for both the coral and *Breviolum* sp. The *P. strigosa* transcriptome-derived Gene Ontologies (GO) were supplemented with GoPredSim ProtT5 protein language model ^65^ derived topGO terms using FANTASIA scripts for increased accuracy and resolution in GO annotations (https://github.com/MetazoaPhylogenomicsLab/FANTASIA/tree/main), and tested with classic Fisher’s exact test for enrichment. KEGG Orthology (KO) classifiers from the EggNOG-mapper predicted proteins were used to annotate and identify protein pathways specifically more enriched by time point using both clusterProfiler ^66^ and tested for enrichment using a hypergeometric approach of a Fisher’s Exact test, followed by a Benjamini–Hochberg False Discovery Rate correction (alpha=0.05) with significant terms visualized as dot and alluvial plots.

STAR-Htseq quantitative files were imported to R, where transcripts were filtered out if not present at least once in all 9 samples within each respective time point (12 AM, 6 AM, 12 PM, and 6 PM). Differential expression was quantified using the R package DESeq ^67^. Contrast comparisons were conducted on the DeSeq object using an “LRT” test across all time points. Transcripts with an adjust p-value of 0.01 for *P. strigosa*, and 0.05 for *Breviolum* and a log 2-fold change of 2x for *P. strigosa* and 1.5x for *Breviolum* were considered significantly differentially expressed. Transcripts with rhythmic expression were identified using the R package LimoRhyde ^68^. LimoRhyde quantified rhythmicity using a cosine-sine linear model ^69^ and significance was measured by an F-test across timepoints using two temporal coefficients, followed by empirical Bayes variance shrinkage and Benjamini-Hochberg False Discovery Rate correction. Transcript peak timing (phase, in hours) was calculated from the arctangent of the sine and cosine coefficients (atan2(sine,cosine) x 24/2π), and amplitude was defined as the vector magnitude of these coefficients, representing the peak-to-trough log2 fold change. Transcripts were considered significantly rhythmic if they had a False Discovery Rate ≤0.01 and amplitude of ≥2 log2 fold change. Lastly, plots were constructed using R packages “ggplot2” ^48^, “chordDiagram” ^70^, “ComplexHeatmap” ^71^, and “treemap” ^72^.

## Results

### Gene expression in *Pseudodiploria strigosa* and *Breviolum* sp. through diel cycles

*Pseudodiploria strigosa* exhibited strong diel transcriptomic trends, with the host showing a surge in gene expression at dawn, whereas the symbiont reflected more gene expression similarity through time (Figure 1A-D). Sequencing of 36 host RNA libraries resulted in ∼ 52 million raw reads per sample (∼94% > Q30; GC 42-47%). Post-processing of *P. strigosa* Salmon quantifications yielded a total of 505,116 ‘gene’ transcripts (herein referred to as transcripts), which were filtered to 63,096 for downstream analysis. For *Breviolum* sp., STAR alignment yielded a total of 41,925 transcripts, with 29,526 remaining after filtering.

**Figure 1:**
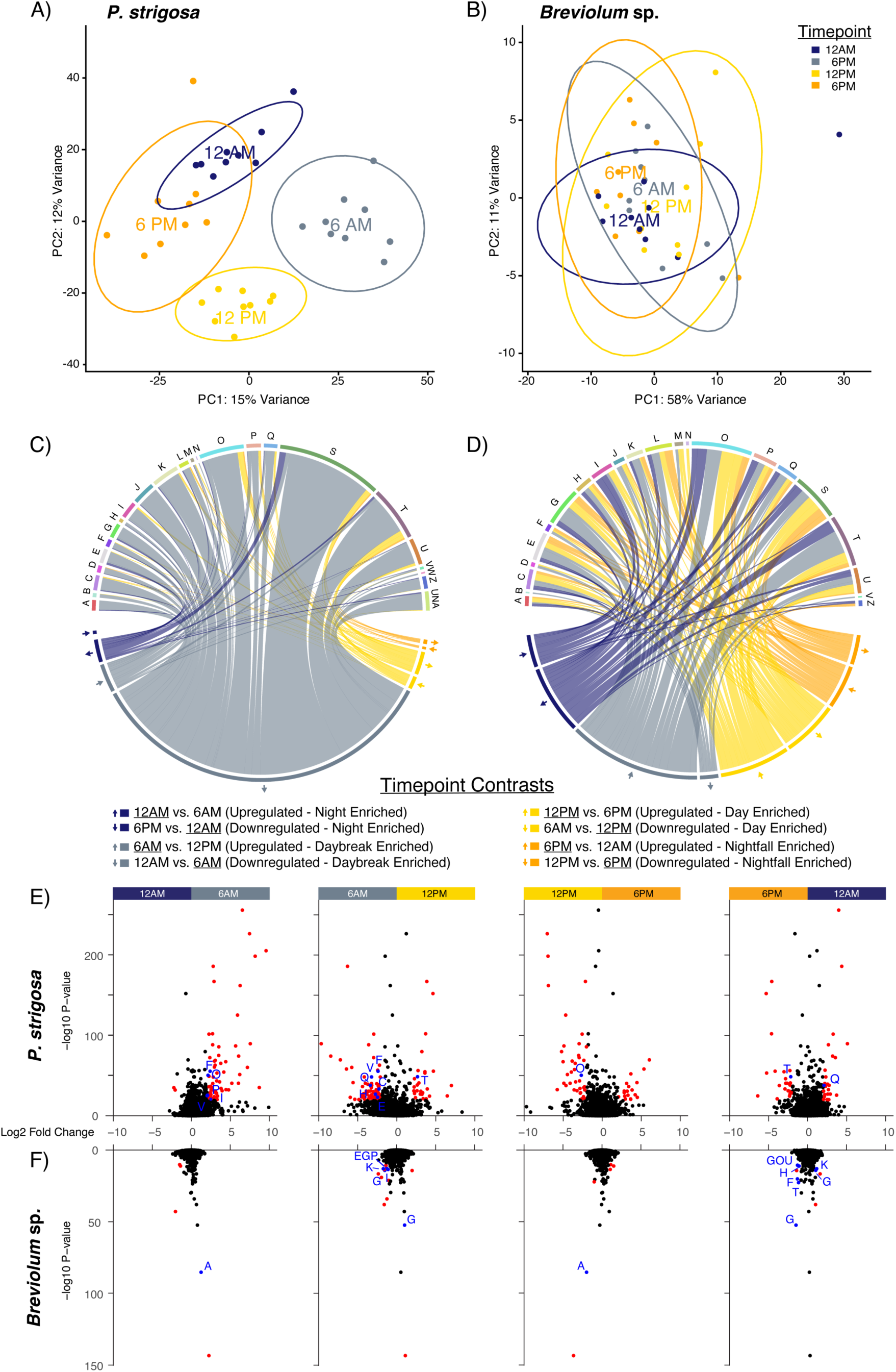
(A-B) Principal components sample scatter plot (A) *Pseudodiploria strigosa* and (B) *Breviolum* sp. *with samples colored for time point* (T12A: midnight, “navy blue”; T6A: dawn, “grey”, T12P: midday, “yellow”; T6P: dusk, “orange”). Each time point was taken in triplicate across three days, totaling n=9 per time of day. (C-D) Chord plots illustrating the transcript contrasts for (C) *P. strigosa* (adjusted p-value < 0.05 and Log2FoldChange = 1) and (D) *Breviolum* sp. (adjusted p-value < 0.05 and Log2FoldChange = 0.58) between specific time points in the order from left to right: Midnight, navy blue; dawn, grey; day, yellow; and dusk, “orange”. Each time point chord connection leads to a specific COG (Clusters of Orthologous Genes) category, A-Z, illustrating the annotated differentially expressed (DE) transcripts based on the contrast between time points, where 1 transcript is 1 COG-annotated DE transcripts, where the thickness represents that number of transcripts. (E-F) Volcano plots showing DE transcripts for *P. strigosa* (E) and *Breviolum* sp. (F) across time contrasts based on log2 fold change on the x-axis and significance using -log10 p-value on the y-axis. Genes with a -log10 p-value greater than 20 and absolute Log2FoldChange 2x (*P*. *strigosa,* Log2FoldChange = 1) and 1.5x (*Breviolum* sp., Log2FoldChange = 0.58) are colored red, and those significant and annotated in blue with corresponding COG categories listed. COG categories are listed as follows: A – RNA processing and modification, B – Chromatin structure and dynamics, C – Energy production and conversion, D – Cell cycle control, division, chromosome partitioning, E – Amino acid transport and metabolism, F – Nucleotide transport and metabolism, G – Carbohydrate transport and metabolism, H – Coenzyme transport and metabolism, I – Lipid transport and metabolism, J – Translation, ribosomal structure, biogenesis, K – Transcription, L – Replication, recombination, repair, M – Cell wall/membrane/envelope biogenesis, N – Cell motility, O – Posttranslational modification, turnover, chaperones, P – Inorganic ion transport and metabolism, Q – Secondary metabolites biosynthesis, transport, S – Function unknown, T – Signal transduction, U – Intracellular trafficking and secretion, V – Defense mechanisms, W – Extracellular structures, Z – Cytoskeleton, and colloquially termed “UNA” – Unassigned.

#### Time-dependent gene dispersion and identifying clusters of orthologous genes

In *Pseudodiploria strigosa*, gene expression showed clear diel structure, with samples clustering primarily by timepoint in principal components analysis (PCA) space. In contrast, its symbiont *Breviolum* sp. displayed weaker temporal organization (Figure 1A&B). Gene dispersion was calculated using variance stabilizing transformation (VST) and plotted in PCA space for both *P. strigosa* (Figure 1A: PC1 = 15% and PC2 = 12%) and *Breviolum* sp. (Figure 1B: PC1 = 58% and PC2 = 11%) to visualize time-specificity.

Dawn represented the transcriptional peak for the host, whereas *Breviolum* sp. exhibited more uniform gene expression through time (Figure 1C&D). DESeq2 contrasts (2x log fold change) revealed 3,693 differentially expressed (DE) genes in *P. strigosa* at dawn, far exceeding midday (305), dusk (50), and midnight (521). Among these, 1,259 DE transcripts were annotated by clusters of orthologous groups (COGs), with 1,166 and 93 mapping to the “dawn vs noon” and “midnight vs dawn” comparisons, respectively (Figure 1C). In contrast, far fewer COG-annotated genes were upregulated at midday (116), dusk (22), or midnight (74), indicating a strong enrichment of functional gene activity at dawn, rather than a gap in annotation. In *Breviolum* sp., fewer DE transcripts were recovered overall; to capture meaningful patterns, contrasts were explored at 1.5x fold change, revealing 160 DE transcripts at dawn compared to midday (145), dusk (102), and midnight (120) (Figure 1D).

In *P. strigosa* dawn was the only timepoint with COG-annotated DE transcripts linked to cell motility (N) and extracellular structures (W), highlighting a unique early-morning transcriptional signature (Figure 1C). While dawn exhibited broad activation across all COG categories, other timepoints showed narrower functional profiles. Midday showed the second largest number of COG-annotated DE transcripts, primarily related to post-translational modification, turnover, chaperones (O), and signal transduction mechanisms (T). Dusk had the fewest COG-annotated DE transcripts overall and was associated with cell cycle control, division, chromosome partitioning (D) and replication, recombination, repair (L). Midnight DE transcripts were primarily associated with post-translational modification, turnover, chaperones (O), signal transduction mechanisms (T), and unknown functions (S), consistent with cellular maintenance and regulatory activity. Volcano plots reflecting pairwise timepoint contrasts (Figure 1E) illustrate that dawn contributes the largest set of unique functional genes. Specifically, dawn DE transcripts encompass five core metabolic categories: energy production and conversion (C), amino acid transport and metabolism (E), nucleotide transport and metabolism (F), lipid transport and metabolism (I), and inorganic ion transport and metabolism (P).

In *Breviolum* sp., COG-annotated DE transcripts showed weaker diel structuring compared to the host, with fewer DE transcripts overall and more subtle time point specificity (Figure 1C&D). Dawn featured unique DE transcripts mapped to chromatin structure and dynamics (B) and defense mechanisms (V), and a pronounced increase in carbohydrate transport and metabolism (G) and signal transduction (T). Uniquely, midday recovered DE transcripts annotated to cell motility (N). Several metabolic categories, including nucleotide transport and metabolism (F) and translation, ribosomal structure, biogenesis (J), were expressed at all time points except midnight. Volcano plot contrasts reflect the limited number of DE transcripts compared to the host (Figure 1F), with dawn and dusk contributing the most unique COG-annotated DE transcripts. Dawn shared three unique COGs with the host, amino acid (E), lipid (I), and inorganic ion (P) transport and metabolism, while dusk contributed unique genes in nucleotide transport and metabolism (F), coenzyme transport and metabolism (H), posttranslational modification, turnover, chaperones (O), and intracellular trafficking and secretion (U).

#### Time-dependent functional gene annotations

Differential expression analysis identified 1,558 significant transcripts in *Pseudodiploria strigosa* and 536 in *Breviolum* sp. (Figure 2A&B). In *P. strigosa,* the heatmap dendrogram revealed two primary sample clusters along the x-axis: one composed predominantly of midnight and dawn samples, and a second containing most midday and dusk samples (Figure 2A). In contrast, *Breviolum* sp. samples exhibit tighter clustering by individual time point, forming more discrete groups (Figure 2B).

**Figure 2:**
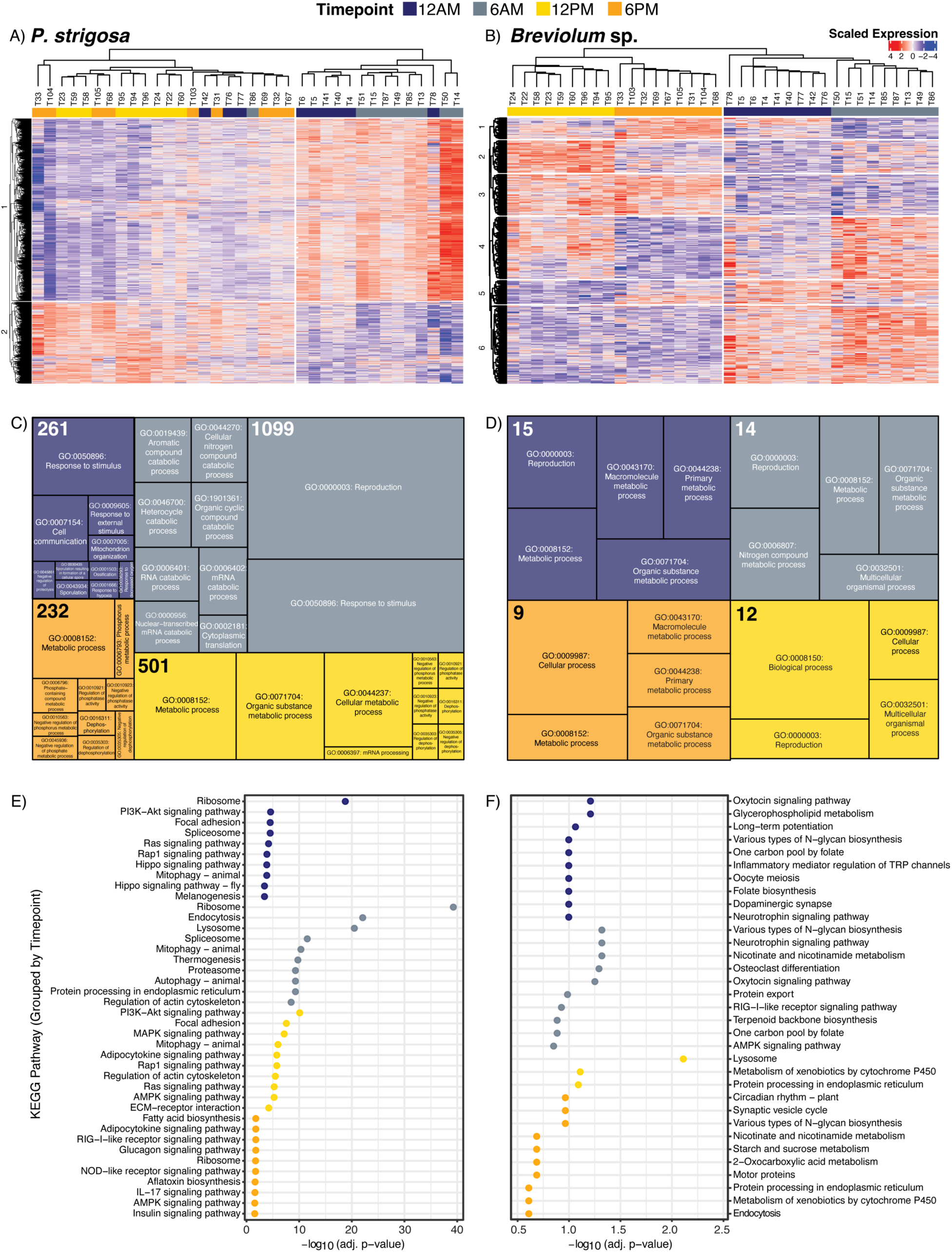
(A-B) Scaled gene expression complex heatmaps of (A) *Pseudodiploria strigosa* and (B) *Breviolum* sp.) with samples clustered using a hierarchical dendrogram showing the expression across the gene set between time points (T12A: midnight, “navy blue”; T6A: dawn, “grey”, T12P: midday, “yellow”; T6P: dusk, “orange”). Significant transcripts were achieved by filtering samples based on adjusted p-value (<=0.01) derived by a Wald test. (C-D) Treemaps showing the top 10 significant GO enrichments for (A) *P. strigosa* and (B) *Breviolum* sp. for each time point, where total significant annotations (bolded number indicated in top left corner of each time point) were calculated through topGO based on a classic Fisher’s Exact test and grouped by time point. E-F) Dotplots showing the top enriched KEGG pathways (n = 10 cutoff) grouped by time point for (E) *P. strigosa* and (F) *Breviolum* sp. and the respective negative log_10_ adjusted p-value. KEGG pathways annotated to “Human Diseases” were filtered out, though can be found in Supplemental Figure 2.

Significant gene ontology (GO) enrichments were identified for differentially expressed transcripts at each timepoint (Figure 2C&D). In *P. strigosa,* dawn exhibited the largest number of significantly enriched terms (1,099), consistent with the high number of DE transcripts at this time point (Figure 2C). Enriched dawn functions primarily involve RNA metabolism and protein turnover, reproduction, stimulus response, and organic compound metabolism. Midday (501 enriched terms) represented several phosphate metabolic and regulatory processes, while dusk (232) shared many of these regulatory mechanisms with lower overall enrichment. Midnight (261) was enriched for hypoxia-related responses, mRNA catabolic processes, mitochondrial organization, and ossification. The photosymbiont *Breviolum* sp. displayed far fewer enriched GO terms at all timepoints (dawn: 23 annotations, midday: 70 annotations, dusk: 19, and midnight: 21), consistent with its lower number of DE transcripts (Figure 2D). Dawn GO enrichments were annotated to immune-related functions and anatomical structure formation, whereas midday was dominated by metabolic processes such as nitrogen compound and heterocycle metabolism. Dusk showed significant enrichment for vesicle-mediated transport accompanied by protein-containing complex assembly and organization. Midnight enrichment was predominantly annotated to cell-cycle and division, and chemical/ionic homeostasis.

In *P. strigosa*, KEGG-annotated DE transcripts were enriched at midnight and dawn with communication, stress signaling cascades (PI3K-Akt, Ras, Rap1, and Hippo) and housekeeping processes such as dealing with damaged cellular components (lysosome, mitophagy, proteasome, and autophagy), and processing new cellular machinery (protein processing, spliceosome, endocytosis) (Figure 2E). Midday DE transcript KEGG enrichment shifted toward growth and proliferation pathways, including PI3K-Atk, MAPK, adipocytokine, Rap1, Ras, AMPL, and insulin signaling, as well as structural remodeling via focal adhesion, regulation of actin cytoskeleton, and ECM-reception interactions. Dusk also exhibited enrichment in energy-related signaling pathways (Adipocytokine and AMPK), with immune-related signaling (RIG-I-like receptor, NOD-like receptor, and IL-17), as well as biosynthetic processes such as fatty acid and glucagon metabolism.

*Breviolum* sp. exhibits a suite of enrichment KEGG terms distinct from the host *P. strigosa,* reflecting temporally structured but non-overlapping transcriptional regulation of the photosymbiont (Figure 2F). Midnight-enriched DE transcripts were significantly enriched for KEGG pathways classically described in neuronal signaling modules such as oxytocin and neurotrophin signaling, dopaminergic synapse, and long-term potentiation. Additional significant enrichment of glycerophospholipid metabolism, folate/one-carbon metabolism, and N-glycan biosynthesis occurred at midnight. At dawn, some of these pathways were also enriched (N-glycan biosynthesis, neurotrophin and oxytocin signaling, folate/one-carbon metabolism), while new enrichments emerged, including protein export, nicotinate and nicotinamide metabolism, RIG-I-like receptor signaling and terpenoid biosynthesis. Midday DE transcripts were predominantly enriched for lysosome activity, xenobiotic metabolism, and protein processing in the endoplasmic reticulum (ER). By dusk, the DE transcripts showed similar enrichment to dawn (nicotinate and nicotinamide metabolism) and midday (xenobiotic metabolism by cytochrome P450 and protein processing in the ER), with new enrichment for circadian rhythms, synaptic vesicle cycle, and energy metabolism pathways (starch, sucrose, and 2-oxocarboxylic acids) (Figure 2F).

Enriched KEGG pathways were mapped across timepoints using narrative Sankey plots (Figure 3A&B), highlighting the differences between the coral host and symbiont in genetic information processing (Figure 3A) and metabolic functions (Figure 3B). In corals, cellular machinery required for transcription and translation, chromosome-related processes, protein folding, sorting, and degradation (Figure 3A) was upregulated from midnight to dawn and largely inactive from midday to dusk. In contrast, *Breviolum* sp. showed no significant enrichment across most timepoints, except at dusk, where the pathways associated with replication and repair, folding, sorting, and degradation were enriched. Transcriptomic signatures in *P. strigosa* showed clear diel shifts in metabolic pathway enrichment (Figure 3B). Midnight and dawn samples were enriched for KEGG pathways related to glycan, xenobiotics, energy, and amino acid metabolism, with dawn showing the most significance to distinct enriched terms. By midday, metabolic pathway enrichment shifted toward lipid and amino acid metabolism, along with “global and overview maps” categories. Dusk samples were enriched primarily for carbohydrate, lipid, non-proteinogenic amino acid, and secondary metabolic pathways. In contrast, *Breviolum* sp. exhibited few significantly enriched metabolic pathways at any timepoint.

**Figure 3:**
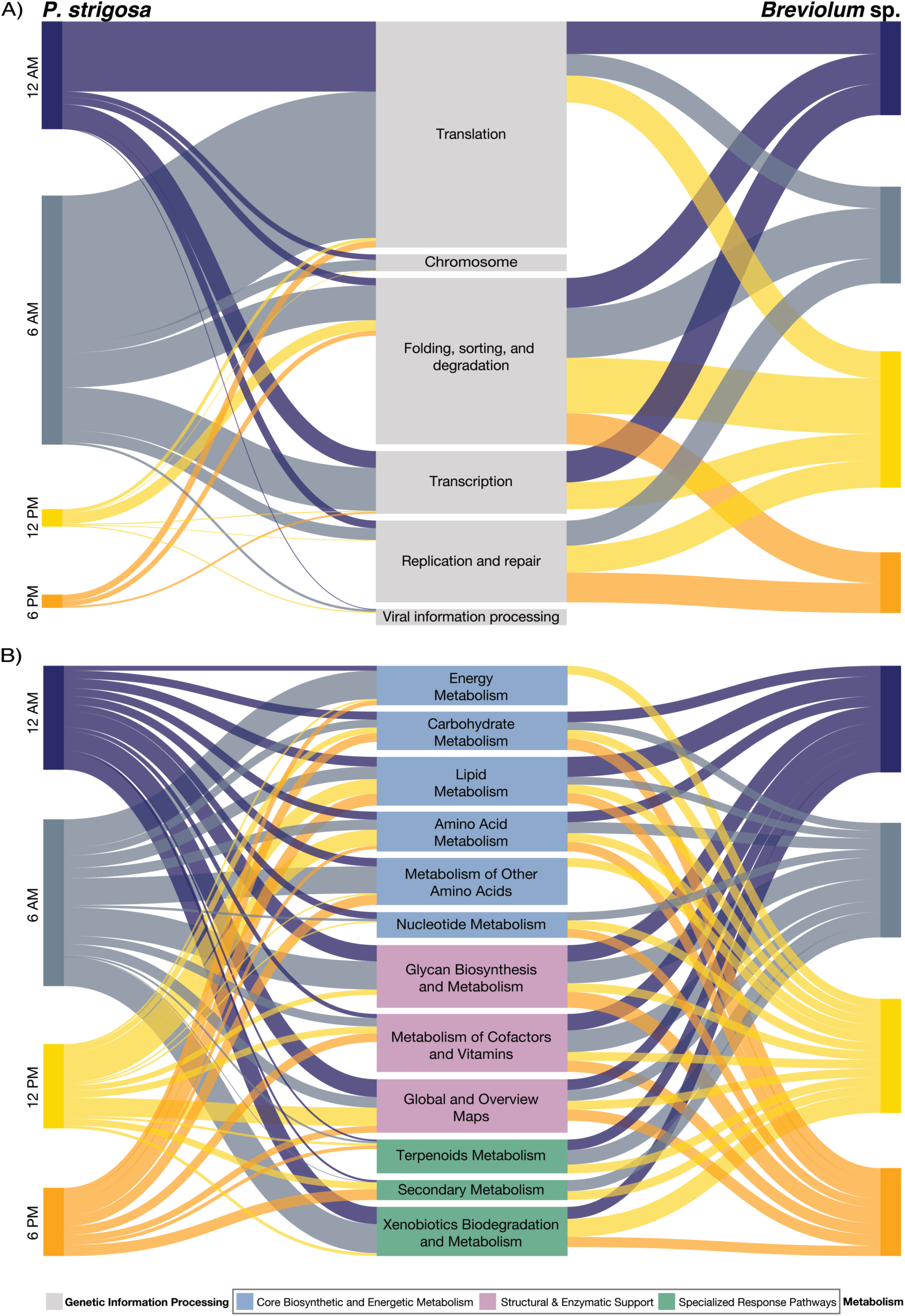
Narrative Sankey plots for KEGG pathways (A) genetic information processing and (B) metabolism for *Pseudodiploria strigosa* (left) and *Breviolum* sp. (right) showing the relationship between time point and the corresponding KEGG pathway. Alluvial line thickness reflects significance calculated using the -log_10_ of the adjusted p-value, where thicker lines indicate higher -log_10_ adj. p-values comparatively. Metabolism is broken into subcategories, grouping core biosynthetic and energetic metabolism (blue), structural and enzymatic support (pink), and specialized response pathways (green).

Significantly rhythmic transcripts from both the coral host and the symbiont were filtered to determine the proportion of transcripts that followed diel periodicity from the total (Figure 4A&B). In *P. strigosa*, 18,833 of 63,096 transcripts met the significance threshold (adjusted p-value <0.01), with only 1,088 being assigned functional annotations (Figure 4A). *Breviolum* sp. displayed 7,999 rhythmic transcripts (of 29,526), with only 71 annotated (Figure 4B). Rhythmic COG-assigned transcripts in *P. strigosa* were dominated by signal transduction, post-translational modifications/protein turnover, and transcription/translation functions (Figure 4A). In contrast, the few rhythmic COG-assigned *Breviolum* sp. transcripts were associated with energy production, carbohydrate metabolism, and protein turnover (Figure 4B).

**Figure 4:**
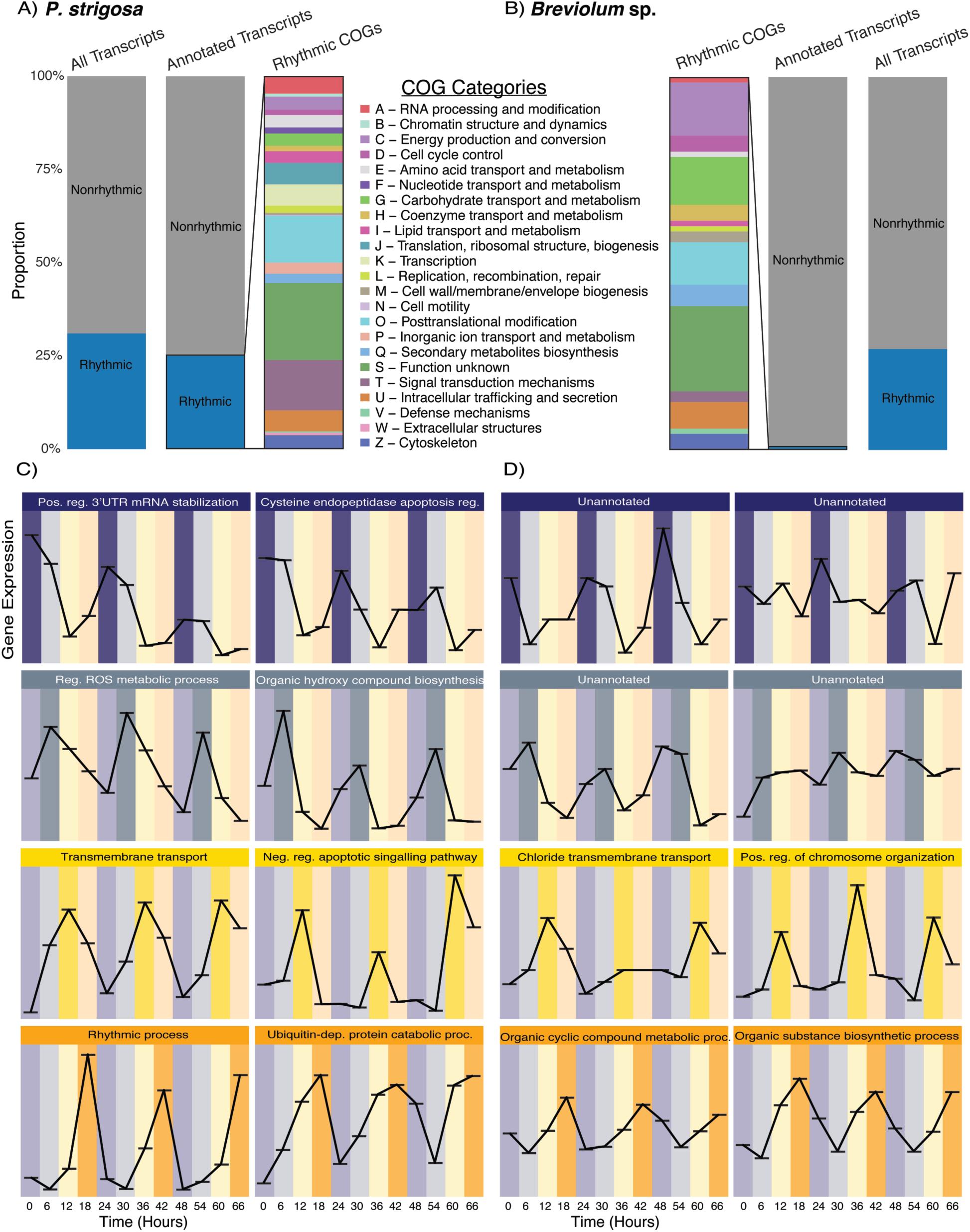
(A-B) Barplots for (A) *P. strigosa* and (B) *Breviolum* sp. illustrating the proportion of total transcripts that are rhythmic vs. non-rhythmic, proportion of annotated transcripts that are rhythmic vs. non-rhythmic, and finally the proportion of rhythmic COG-annotated transcripts. (C-D) Line graphs visualizing significantly rhythmic transcripts recovered from (C) *Pseudodiploria strigosa* and (D) *Breviolum* sp. across three diel cycles. The x-axis represents the hours over the three days, and the y-axis represents transcript expression in transcripts per million (TPM) for *P. strigosa* and raw counts for *Breviolum* sp. The line graphs are separated by time points wherein the transcripts are specific in peak to trough amplitude: midnight, “navy blue”; dawn, “grey”, midday, “yellow”; and dusk, “orange”. Transcripts were annotated for their highest gene ontology assignment resolution. Corresponding quantitative boxplots can be found in the supplementary section for each transcript in Supplementary Figure 3.

To visualize clock-controlled gene (CCG) transcript expression through time, representative rhythmic transcripts are shown for each timepoint for both *P. strigosa* and *Breviolum* sp. (Figure 4C&D). In *P. strigosa*, midnight CCG transcripts included regulators of mRNA stability and cellular control through apoptosis (Figure 4C). Dawn featured CCG transcripts associated with reactive oxygen species (ROS) mitigation and hydroxy compound biosynthesis. Midday *P. strigosa* transcripts were enriched for transmembrane transport and apoptotic suppression. Transcripts with expression peaking at dusk highlighted functions related to rhythmic process and ubiquitin-dependent protein catabolism. In the symbiont *Breviolum* sp., far fewer rhythmic transcripts were annotated, resulting in unannotated representative CCG transcripts for both midnight and dawn (Figure 4D). For the CCG transcripts that were annotated, midday featured chloride transmembrane transport and chromosome organization, while transcripts peaking at dusk were enriched for cyclic-compound and organic-substance metabolism.

### Pseudodiploria strigosa microbial assemblages through diel cycles

To explore how the *Pseudodiploria strigosa* microbiome responds to diel cycles, both prokaryotic (16S rRNA) and eukaryotic (18S rRNA) genes were amplified at the V4 region and sequenced from 36 coral and 12 seawater samples. A total of ∼23.2 million 16S and ∼25.0 million 18S raw reads were recovered and subsequently quality filtered resulting in ∼20.3 million and ∼13.1 million processed reads, respectively. After removing spurious taxa, the amplicon sequence variant (ASV) table comprised 6,521 prokaryotic and 1,506 microeukaryotic ASVs, including 588 Symbiodiniaceae ASVs across all samples.

#### Microbial stability across the holobiont

Alpha diversity was quantified using the Shannon index for diversity and observed richness partitioned by prokaryome, eukaryome (excluding Symbiodiniaceae, Metazoa, and Embryophyceae), and Symbiodiniaceae (Figure 5A-F). Overall, the prokaryome exhibited the highest diversity and richness, whereas the eukaryome was less diverse but comparatively more stable across timepoints (Figure 5A-F). Symbiodiniaceae showed low diversity and richness yet remained the most temporally stable of the three communities. Within the prokaryome, Shannon diversity peaked at dawn and declined toward dusk, with mean Shannon values between 3 and 4 across timepoints, displaying apparent diel periodicity (Figure 5A). Observed richness mirrored this pattern, with a notable increase in variability at dawn, ranging between 750 and 1250 taxa, with the highest variability at dawn and lowest variability at midnight (Figure 5D).

**Figure 5:**
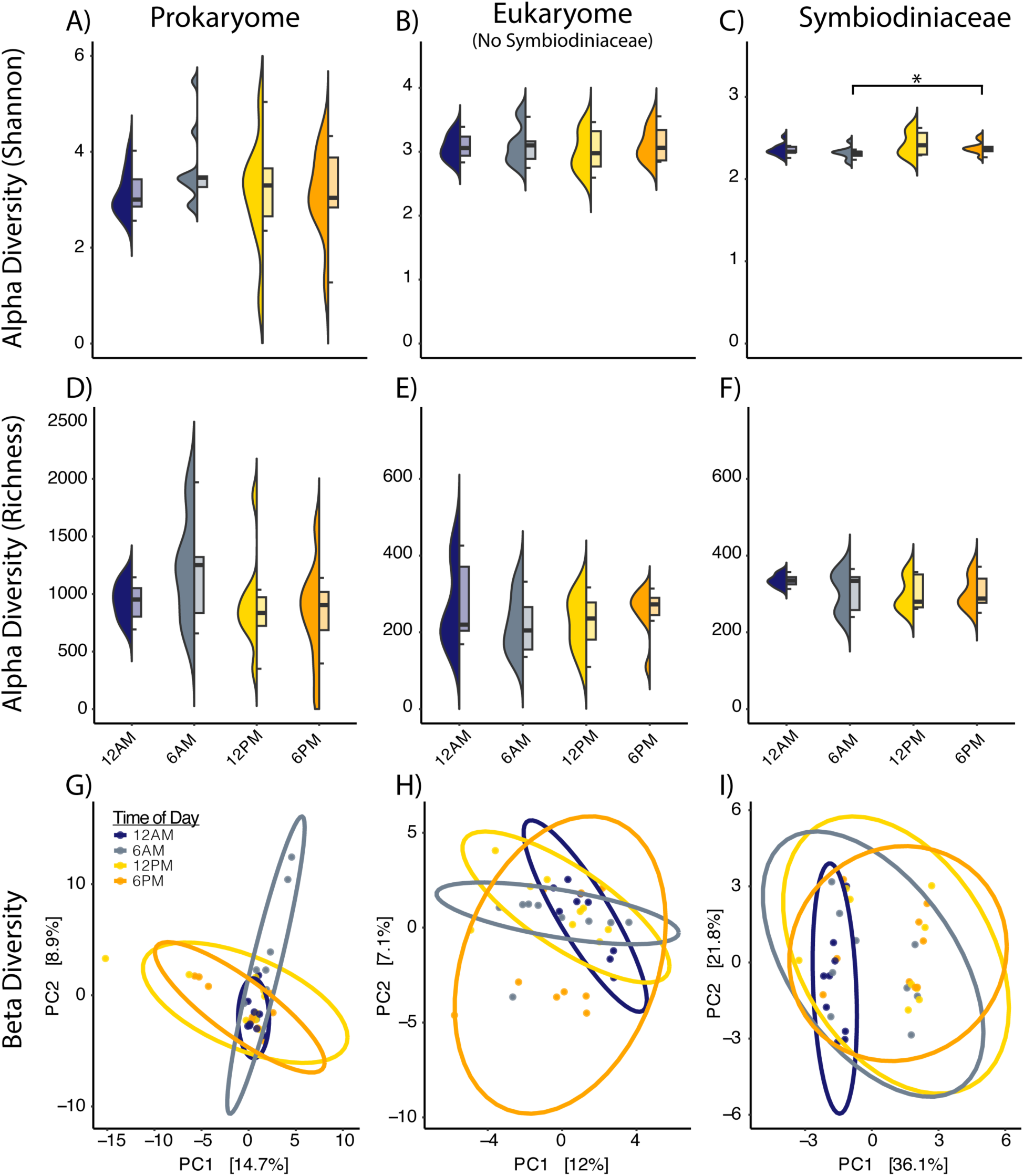
*Pseudodiploria strigosa* microbial alpha and beta diversity plots showing Shannon diversity (A-C), observed richness (D-F), and Aitchison distance PCA (G-I). Samples are grouped by time points: midnight (12 AM), dawn (6 AM), midday (12 PM), and dusk (6 PM). Plots are separated by their kingdom, Prokaryota (A, D, G), Eukaryota without Symbiodiniaceae (B, E, H), and finally Symbiodiniaceae (C, F, I). Asterisk indicates significance by Wilcoxon rank-sum p<0.05. Significance in beta clusters were tested using adonis2 PERMANOVA.

The average eukaryotic Shannon diversity shows significantly less variability than the prokaryome, with mean values between 2.75 and 3.25 across timepoints and only subtle diel trends (Figure 5B). Eukaryotic richness peaked at dusk, then declined at midnight and dawn, and rose at midday (Figure 5E). Symbiodiniaceae showed minimal variation, with Shannon diversity values between 2.25 and 2.5 across timepoints, and the least variability at midnight (Figure 5C&F). Pairwise testing revealed that Shannon diversity differed significantly (Wilcoxon rank-sum, p<0.05) only from dawn to dusk; observed richness was higher at midnight and dawn than during the daylit timepoints.

Beta diversity of *P. strigosa* microbial assemblages showed weak but significant diel structuring by timepoint, with sampling day included in the model to account for potential confounding effects (Figure 5G-I). Prokaryotic communities were strongly influenced (PERMANOVA R^2^=0.65, 0=0.003), while eukaryotic assemblages, including Symbiodiniaceae, show weaker but significant effects (PERMANOVA R^2^=0.11 and 0.18, 0=0.014 and 0.009, respectively). Coral microbial diversity and community compositions were different from the surrounding seawater (Supplementary Figure 4A-H), where Shannon diversity indices were ∼5.0 and ∼6.0, and richness averaged ∼2000 and ∼3000 for prokaryotic and eukaryotic communities, respectively. In contrast, seawater beta diversity plots were primarily structured by sampling day rather than timepoint, with prokaryotes (PERMANOVA R^2^=0.47, 0=0.003) and eukaryotes (PERMANOVA R^2^=0.38, 0=0.002) displaying day-driven clustering (Supplementary Figure 4C&F).

#### Microbial community composition and highly abundant taxa

Prokaryotic microbial composition in *Pseudodiploria strigosa* weas highly stable across the diel cycle, with only a subset of taxa exceeding 0.1% relative abundance (Figure 6A). The prokaryotic fraction of the microbiome was dominated by *Endozoicomonas*, *Listeria*, *Bacteroides*, *Stenotrophomonas*, *Pseudomonas*, *Ralstonia*, *Ruminococcus* (torques group), *Flavobacterium*, and *Staphylococcus*, with only minor diel fluctuations. Subtle shifts were observed in *Ralstonia*, *Delftia*, *Pseudomonas*, *Bacteroides*, and *Endozoicomonas*, where *Endozoicomonas* decreased slightly during daytime hours, in constrast to the other aforementioned taxa that showed slight reductions at night. Archaea were low in relative abundance but compositionally stable when analyzed separately, dominated by Marine Group II and III (orders), *Nitrosotalea* (genus), Bathyarchaeia (class), Woesearchaeles (order), *Nitrosopumilus*, *Nitrosophelagicus*, *Nitrocosmicus, Nitrosophaera*, *Nitrosotenuis*, and *Cenarchaeum* (Figure 6B).

**Figure 6:**
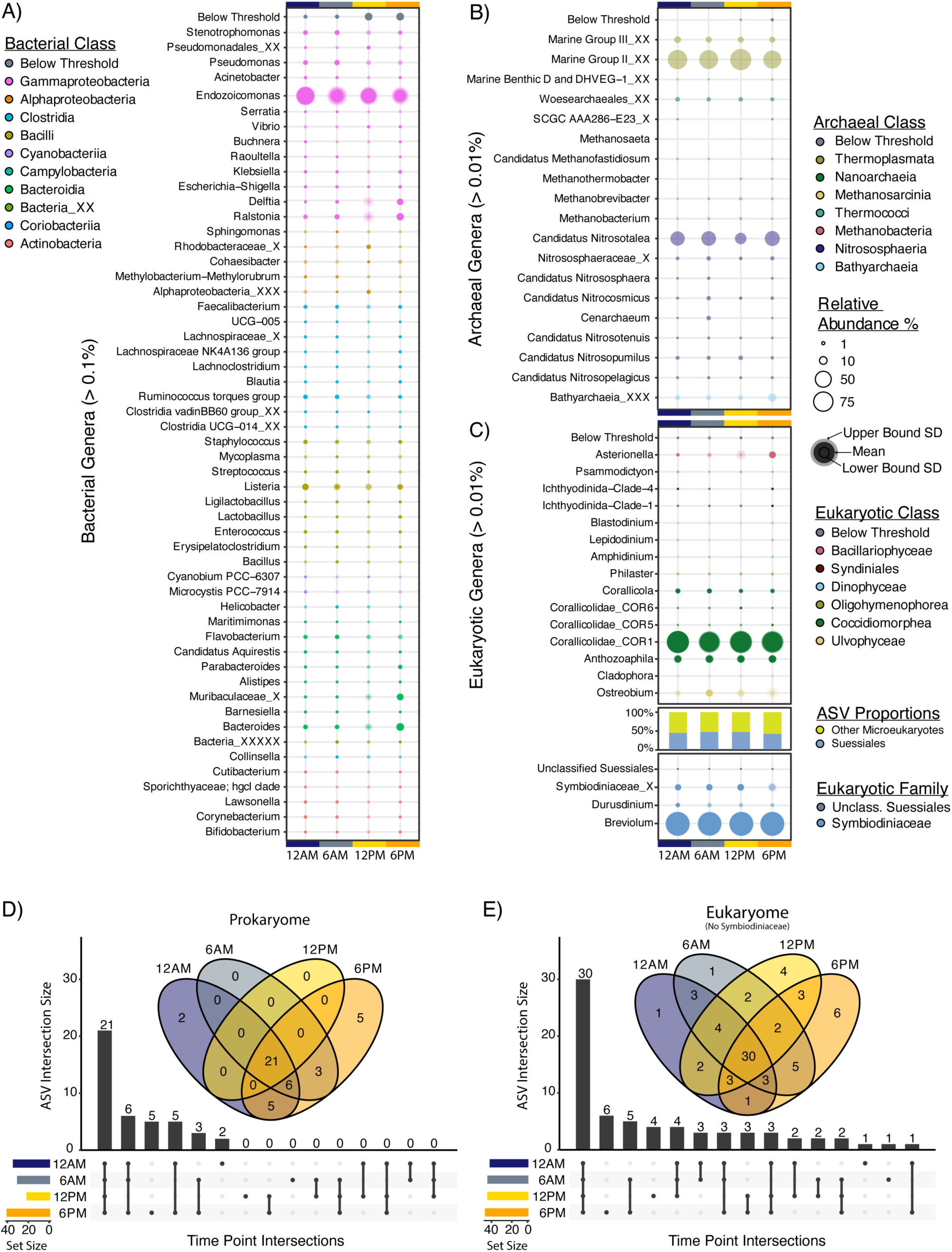
Bubble plot showing phylogenetically sorted taxa for (A) the bacterial fraction, (B) the archaeal fraction, and (C) the eukaryotic fraction at the genus level community composition on the y-axes by time of day (12 AM, 6 AM, 12 PM, 6 PM) on the x-axis. The color of the bubble indicates the taxonomic class. The size of the bubble indicates the average relative abundance by percent. In contrast, the lighter shade of the bubble represents the upper bound of the standard deviation, and the darkest shade represents the lower bound of the standard deviation. The “Below Threshold” category is the summation of the remaining taxa that do not contribute greater than 0.1% (bacterial) 0.01% (archaeal), and 0.01% (eukaryotes) relative abundance. The “X”s following the taxonomy denote taxonomic placement from the highest taxonomic classification. Eukaryotic reads are separated by the class Suessiales and the remainder of the microeukaryotic fraction. Core microbiome of *P. strigosa*, visualized by microbial domain, (D) prokaryome and (E) eukaryome (excluding Symbiodiniaceae) consisting of a Venn diagram showing the overlapping core ASV taxa by time point (midnight: T12A, dawn: T6A, midday: T12P, and dusk: T6P), respectively, and upset plot detailing the intersection (and ASV set size) of each time point and the unique ASVs accordingly sorted by descending frequency. The core microbiome is calculated using a core detection of 0.0001 (0.1%) and a prevalence of 0.75 (in at least 75% of samples).

The microeukaryotic community of *P. strigosa* was dominated by members of the family Symbiodiniaceae (Class: Suessiales), primarily *Breviolum* with a secondary association to *Durusdinium* (Figure 6C). *Durusdinium* increased in relative abundance in the midnight hours before declining through the day, whereas the overall proportion of ASVs assigned to Suessiales remained relatively stable (44 - 47% across timepoints). The proportion of Suessiales within the coral host is comparatively much larger than the proportion in the surrounding seawater with 5-15% (Supplementary Figure 4H). Beyond Suessiales, the most prevalent microeukaryotes belong to the family Corallicolidae (Apicomplexa), *including Corallicola*, *Anthozoaphila,* and Corallicolidae COR1, COR5, and COR6 (Figure 6C). Other frequent taxa included the endolithic photosynthetic alga *Ostreobium,* diatoms (*Asterionella, Psammodictyon)*, Syndiniales (Ichthyodinida), several Dinophyceae, and the scuticociliate *Philaster* (Figure 6C). Over three days of sampling, the *P. strigosa* microbiome exhibited remarkable temporal stability, with both prokaryotic and eukaryotic dominant community members showing minimal diel turnover (Supplemental Figure 6A&B). *Endozoicomonas* dominated the prokaryome and remained consistently abundant in every sample, highlighting its role as a persistent core symbiont (Supplementary Figure 6A). In contrast, few taxa showed consistent diel fluctuation; *Ostreobium* displayed consistent decrease at midnight, while other microeukaryotic taxa including those from Corallicolidae exhibited high diel stability across all samples across all timepoints (Supplementary Figure 6B).

#### Core microbiome and rare taxa

Core microbial taxa were recovered using detection of 0.1% and presence in at least 75% of samples (Figure 6 D&E). Within the prokaryotic microbiome, 21 core ASVs were shared across all time points. Only midnight and dusk harbored unique ASVs: *Raoultella* sp. (ASV66) and *Stenotrophomonas* sp. (ASV529) were specific to midnight, whereas dusk hosted *Blautia* sp. (ASV151), *Mycoplasma* sp. (ASV365), *Streptococcus* sp. (ASV446), *Maritimimonas* sp. (ASV670), and *Stenotrophomonas* sp. (ASV761) (Figure 6D). Dusk exhibited the largest intersection set, sharing the most ASVs with other timepoints, whereas midday displayed the smallest, with few unique or overlapping ASVs.

Core eukaryotic microbial taxa in *P. strigosa* were highly conserved across diel cycles, with only a fraction of ASVs unique to specific timepoints (Figure 6B). The core community consisted of 30 ASVs shared across all timepoints, dominated almost entirely by corallicolids (25 COR1 and 4 *Anthozoaphila* ASVs), and a single *Ostreobium* sp. (ASV449). Unique ASVs were rare and showed subtle temporal biases: midnight and dawn each harbored a single COR1 ASV (ASV531 and ASV647), midday added four mostly corallicolid ASVs, and dusk contributed six ASVs enriched in *Ostreobium* and *Anthozoaphila*. This pattern highlights a striking temporal stability in the eukaryotic microbiome, with dusk exhibiting diverse core taxa.

#### Differentially abundant and co-occurring microbial taxa in the P. strigosa microbiome

Despite the broad temporal stability of the *P. strigosa* microbiome, subsets of microbial taxa exhibit diel specificity (Supplementary Figure 7A-D). Prokaryotic taxa dominated these shifts: day-enriched taxa include *Hoeflea sp.*, *Butyricicoccus sp*., *Polaromonas sp*., *Pseudohongiella sp*., *Tropicimonas sp.*, *Parabacteroides sp*., *Hyphomonas sp*., *Actibacterium sp*., *Cohaesibacter sp*., *Clostridium colicanis*, and *Cribrihabitans sp.*, whereas night-enriched taxa were limited to *Scytonema sp.* (VB-61278), *Cupriavidus metallidurans*, and *Labrenzia alexandrii* (Supplementary Figure 7A). Eukaryotic differentially abundant taxa were even fewer, largely driven by the diatom *Asterionella glacialis* during the day and corallicolids (*Corallicola* and *Anthozoaphila*) at night (Supplementary Figure 7B). Pairwise comparisons revealed that only intervals 12 hours apart recovered differential taxa, with midday-to-midnight shifts in prokaryotes Lachnospiraceae, *Staphylococcus sp.*, and *Flavonifractor sp*., and dawn-to-duck transitions highlighting SAR324 (marine group B), as well as taxa from Bacteroidia and Entomoplasmatales (Supplementary Figure 7C&D). Together these patterns suggest that only a fraction of the microbiome shows any variation over diel timeframes, supporting the conclusion that the majority of the microbiome is highly stable.

Co-occurrence network analysis demonstrated that the subtle diel microbiome variation is effectively captured through inference association networks (Figure 7A&B). *Breviolum* consistently exhibited the strongest correlation with COR1 corallicolids across all timepoints, suggesting algal-apicomplexan associations. However, day-night variation emerges in the networks: *Breviolum* features correlations with *Stenotrophomonas* and Pseudomonadales during the daytime hours, including unique taxa *Blautia, Defluviicoccus, Hahella*, *Parabacteroides*. Nighttime networks exhibited stronger associations with *Endozoicomonas,* while recovering new associations with *Ostreobium*, *Staphylococcus*, *Streptococcus,* Ruminococcaceae, and methylotrophs (*Methylobacterium-Metholorubrum*). Together, these diel-influenced associations suggest that the otherwise stable coral microbiome undergoes subtle reconfigurations between day and night that likely have functional consequences.

**Figure 7:**
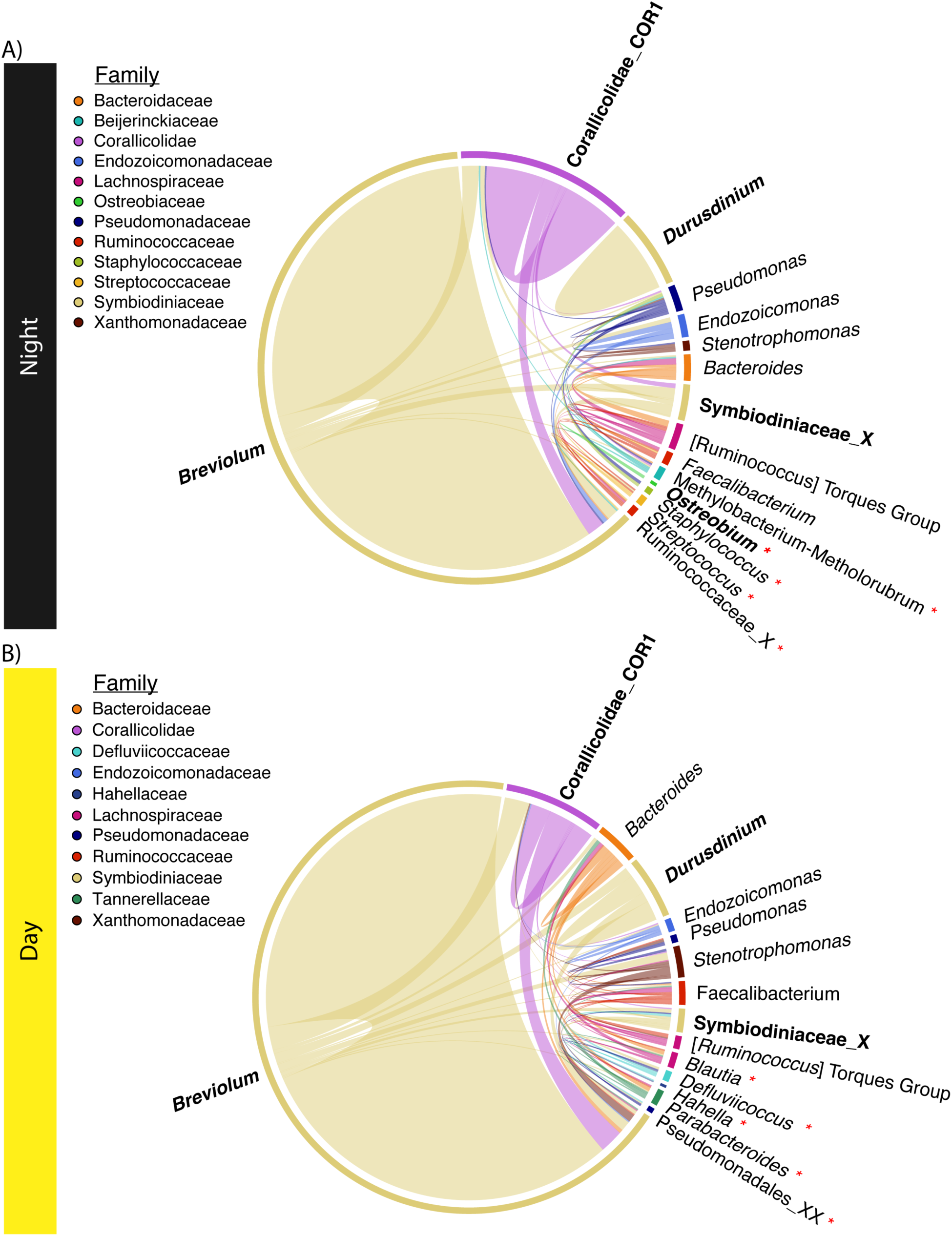
Microbial co-occurrence networks for *P. strigosa* illustrating the significantly correlated associations between prokaryotic (regular text) and eukaryotic (bolded text) genera. Networks compare night and day samples to observe changes in co-occurrence between light and dark periods. Taxa with a red asterisk next to it indicate taxa that shift between day and night. Correlation network method “SpiecEasi” with p-value threshold 0.01 and p-adjusted using false discovery rate.

## Discussion

Most tropical reef-building corals rely on fixed carbon supplied by their resident photosymbionts (in the Family Symbiodiniaceae), which can translocate ∼80% of the coral’s daily carbon demand as photosynthates ^73^. In *Pseudodiploria strigosa,* the dominant symbionts in the genus *Breviolum* fuel host metabolism, driving calcification, growth, and repair. This intimate symbiotic relationship has evolved to be a critical driver of the success of most shallow-water tropical and subtropical corals. Because this light-driven carbon flux is vital to the host on a 24-hour cycle, we profiled the coral and symbionts every six hours over three consecutive days. The approach revealed pronounced diel periodicity in the host, while *Breviolum* symbionts showed more subtle, phase-shifted transcriptomic changes. In contrast, the microbiome community remained largely stable, with minor genus-level fluctuations reflecting light and dark periods.

### Gene expression in the coral host and symbiont through time

*P. strigosa* exhibits clear time-specific cellular functions, consistent with prior evidence that early morning environmental light cues trigger key diel responses in corals. Dawn represents a critical transition, marked by a surge in transcript upregulation, driving preparatory functions to withstand the upcoming influx of photosynthates, reactive oxygen species (ROS), and ultraviolet (UV) radiation ^74,75^. Our results support this notion, as significant COGs recovered from the contrasts show dawn being critical to developing the proteins necessary in energy production and conversion, as well as nucleotide, amino acid, lipid and inorganic ion transport and metabolism. Although early morning light-sensing pathways, such as cryptochromes ^26^ and G-protein-coupled receptors ^76,77^ likely initiate key diel responses, *Breviolum* sp. shows no corresponding surge of transcript expression at dawn. Despite this, the host and its symbiont *Breviolum* sp. upregulate similar processes at dawn, as reflected in the COG annotations, including amino acid, lipid, and inorganic ion transport and metabolism (E, I, and P).

The predominance of upregulated transcripts at dawn in *P. strigosa* coincides with processes typical of cellular/protein turnover and renewal, consistent with the onset of morning environmental conditions. *Breviolum* sp. featured Gene Ontology (GO) enrichment for processes linked to environmental sensing and early-day cellular reorganization potentially for environmental cues and preparatory organelle development (GO: response to external stimulus, immune effector process, small molecule biosynthetic process, anatomical structure formation involved in morphogenesis). This notion is also supported in dawn KEGG pathways relating to housekeeping processes, such as the catabolism of damaged cells (lysosome, mitophagy, proteasome, and autophagy) and the biosynthesis of new cellular machinery (protein processing, spliceosome, and endocytosis). Collectively, the *P. strigosa* data exemplify dawn as a critical molecular trigger for initiating protein biosynthesis, cellular reorganization, and anticipatory mechanisms for daytime environmental stressors, which warrants future validation at the protein level.

Enrichment of phosphate-related processes, including “regulation of phosphatase activity” and “dephosphorylation” at midday, hints at temporal modulation of phosphorylation-dependent signaling and energy turnover. While we cannot ascertain the precise pathway components from transcriptomic data alone, these enriched processes are broadly consistent with cellular remodeling and energy conversion during peak daylight hours ^74^. Contextually, phosphate uptake and turnover are critical for maintaining symbiont-to-host cell balance and supporting carbon translocation under thermal stress ^78^. Functional enrichment of growth-related KEGG pathways, including PI3K-Akt, Adipocytokine, Rap1, Ras, and AMPK, focal adhesion, and extracellular matrix (ECM)-receptor interactions, further points to midday as a period of increased tissue growth and maintenance ^74^. This pattern aligns with previous observations that disruption of ECM structural genes, such as Collagen α-1(II), compromises calcification and cell adhesion under thermal stress ^79^. Although these data do not directly measure phosphate uptake or metabolic flux, the enrichment of energy metabolism pathways (lipid and amino acid metabolism) suggests that midday may represent a phase of active growth and structural development in the host.

Diel oscillations in marine primary production drive carbon fluxes that support marine food webs ^80^, and previous transcriptomic studies have revealed that genes involved in central metabolism, cell cycle progression, and photosynthesis exhibit diel periodicity ^81–86^. Maas et al. (2024) demonstrated that 96% of transcripts in the copepod *Pleuromamma xiphias* transcriptome are differentially upregulated at midday. In our dataset, midday gene expression in *Breviolum* sp. is enriched for lysosomal activity, xenobiotic metabolism, and ER protein processing, consistent with photosynthesis-induced ROS and UV stress accumulation. These enriched processes likely reflect cellular quality control, including degradation and removal of UV-damaged organelles, and enzymatic detoxification of ROS through cytochrome P450. Corals and their symbionts are known to regulate superoxide dismutase and catalase enzymatic activity in response to increased solar UV radiation, as a protective mechanism to support the symbiont through increased ROS production during the day ^87^, and our midday enrichment data support analogous stress-mitigating mechanisms. This aligns with prior observations that algae are hyper-producing carbon during the daylit hours, generating a hyperoxic and carbon-rich surface mucosal boundary layer ^17,88,89^, driving the need for increased detoxification and organelle turnover. Gene ontology enrichment in *Breviolum* sp. for nitrogen-compound and heterocycle-related metabolic processes at midday likely reflects biosynthetic pathways that facilitate macromolecule repair and turnover, rather than direct carbon metabolism ^90^. Additionally, the enrichment of chloride transmembrane transport and positive regulation of chromosome organization among rhythmic midday transcripts suggest roles in ion homeostasis for pH regulation and facilitating access to genes required for macromolecular repair ^74,91,92^. Collectively, these data point to midday as a phase of cellular maintenance in *Breviolum* sp., coordinating metabolic resource allocation and repair processes under peak light stress.

Dusk in *P. strigosa* shows the least consistent gene expression across days, suggesting this timepoint represents a fine-tuning phase of daytime metabolic functions (e.g. phosphatase activity, dephosphorylation, and phosphorous metabolism), while preparing for low-light conditions and reduced photosynthate translocation from *Breviolum* sp. The coral host simultaneously shifts towards immune-related signaling pathways (e.g., NOD-like receptor, IL-17, and RIG-I-like receptor), potentially as a defense mechanism against opportunistic microbes during the transition to low-oxygen conditions ^93^. With the onset of night, the hyperoxic holobiont microenvironment becomes anoxic as its aerobic members continue to respire ^17,88,94^. Despite oxygen depletion, glycolysis can continue anaerobically, while additional carbon and other essential byproducts are likely acquired through heterotrophic feeding ^95^. Unlike the surge in upregulation observed at dawn, dusk is marked by down-modulation of genetic information processing and metabolic functions, consistent with a preparatory phase for cellular repair and recovery. This is supported by the rhythmic transcript expression patterns, with clock-controlled genes peaking at dusk and housekeeping genes involved in ubiquitin-dependent protein catabolism, showing rhythmicity, tagging misfolded or UV-damaged proteins for degradation. Meanwhile, *Breviolum* sp. exhibits upregulation in metabolic processes involving starch, sucrose, and 2-oxocarboxylic acids, likely driven by circadian entrainment by light-sensing pathways ^96^. Gene Ontology terms also indicate enrichment of monoatomic-anion and vesicle-mediated transport, suggestive of photosynthate translocation in the coral host. This pattern is consistent with evidence that dinoflagellate protein metabolism and synthesis are governed by circadian rhythms, with peak activity occurring at dusk ^74,96^.

Midnight shows the fewest differentially expressed transcripts in *P. strigosa*, though it is characterized by unique functional enrichments. KEGG pathways highlight stress-associated signaling cascades (Ras, Rap1, PI3K-Akt, and Hippo), likely reflecting the coral’s response to progressive hypoxia during the dark hours ^74,88,89^. These functional shifts likely prepare the coral for the pronounced abiotic shifts that accompany dawn, when photosynthesis resumes and oxygen levels replenish. KEGG terms at midnight include cell communication and vesicle-associated processes. These map onto “neural-associated” KEGG pathways (e.g., oxytocin and neurotrophin signaling, dopaminergic synapse, and long-term potentiation), which likely reflect shared molecular machinery such as vesicle-mediated trafficking and ion transport. Metabolite exchange may be facilitated by such vesicle-mediated processes between host and symbiont during dark hour stress ^97^, a form of functional crosstalk also observed during thermal stress ^98^. Finally, *P. strigosa* KEGG enrichment shows upregulation of translation, transcription, replication, and repair machinery at midnight, indicating active macromolecule maintenance under low-carbon conditions. In parallel, the coral likely relies on amino acid catabolism to sustain energy production during low levels of photosynthates ^89,99^. Similar night enrichment of DNA repair, RNA biosynthesis, and response to hypoxia has been observed in *Euphyllia paradivisa* ^25^, consistent with circadian regulation of cellular maintenance in corals at night.

These late-night mechanisms likely reflect the maintenance of core housekeeping processes, such as degradation of hypoxia-induced and damaged proteins, and the replenishment of cellular components needed for the surge in transcriptional regulation at dawn. In *P. strigosa*, these patterns indicate a period of reduced metabolic complexity and enhanced cellular maintenance, functionally analogous to “rest” phases described in other invertebrates ^6–9^, though corals may still engage in heterotrophic feeding at night. This pattern represents a temporally regulated state focused on cellular repair and homeostasis during the low-oxygen nighttime period. While few canonical clock genes (e.g., *cry, clock, cycle*) were recovered in our *de novo* transcriptome, this likely reflects annotation limitations rather than their absence. We observe multiple transcripts with strong rhythmic expression that are putative clock-controlled genes (CCGs), inferred from their diel periodicity and functional categories aligned with homeostatic and metabolic cycling. These include genes involved in protein metabolism and turnover, ion transport, and stress mitigation, hallmarks of downstream clock output even when core transcription-translation loops are not fully captured in the data. Future studies leveraging comprehensive genomic annotation and functional assays, such as using CRISPR to knock out key CCGs ^100^, will be critical to verify their regulatory structure and directly link these genes to ecologically critical processes, such as environmental cues involved in synchronized spawning ^26^.

### The stability of the coral microbiome through diel cycles

The coral host *P. strigosa* exhibits pronounced circadian rhythms in gene expression, while its microbiome (bacteria, archaea, protists, fungi, and algae) remains stable across diel cycles. Coral microbiomes have been described in the literature as either stable or variable depending on the host species and health status ^19,101^. However, corals and reef ecosystems experience extreme short-term abiotic fluctuations, including swings in oxygen availability, salinity, sedimentation and turbidity, pH, and temperature ^17,33,88,102–105^. A substantial fraction of the coral-associated microorganisms support homeostatic functions, including the production of antimicrobials (e.g., tropodithietic acid for bacterial suppression) and B vitamin provisioning (e.g., thiamin, biotin, riboflavin), nutrient transfer, and biogeochemical cycling such as photosynthesis and carbohydrate metabolism, and sulfur-cycle processes like dimethylsulfoniopropionate (DMSP) cleavage to dimethyl sulfide ^13,14,106–108^. In this study, *P. strigosa*’s microbiome exhibited subtle diel fluctuations in community composition, as supported by the PERMANOVA and ordination analyses yet the overall structure appears generally stable across three replicated 24-hour cycles. This contrasts with the stochastic temporal variability observed by Caughman et al. (2021), as they reported no clear patterns of periodicity through time, likely due to sparse replication in their sampling design. Species-specific patterns of microbiome stability have been noted previously in corals ^109,110^, and our findings largely support this notion in *P. strigosa*, with subtle diel shifts in the relative abundance of a few taxa between day and night. This is further supported by the findings from Seiblitz et al. (2025) where four species of coral (two zooxanthellate: *Madracis decactis* and *Mussismilia hispida* and two azooxanthellate: *Tubastraea coccinea* and *T. tagusensis*) did not show significant microbiome changes over diel cycles.

This study provides a unique perspective on the coral microbiome as the first study to examine the eukaryotic fraction across diel cycles. Much less focus has been devoted to studying the eukaryotic fraction of the coral microbiome ^13,14^; however, previous literature supports that microeukaryotes host their own microbiomes ^111–114^. Microeukaryotes that feed heterotrophically on bacteria and archaea contribute to the diel turnover of microbial composition ^115^. Diel fluctuations in environmental parameters, such as light, oxygen, and nutrient availability, likely further facilitating microbial restructuring, particularly among Bacteria. In contrast, Symbiodiniaceae exhibit inherently low cellular division rates ^116^, which may underly the minimal composition shifts observed in these communities. However, host-algal interactions, including heterotrophic feeding ^116^ during night hours and midday symbiont expulsion ^117^ to mitigate excess ROS production, still contribute to subtle holobiont-level dynamics. *P. strigosa* primarily associates with *Breviolum* spp., and our data support that finding, with minor co-associations with *Durusdinium trenchii*. Additional taxa, such as the apicomplexan corallicolids have been documented in corals worldwide ^118^, in some cases as abundant as Symbiodiniaceae ^14,119^. Although the functional role of these corallicolids remains unknown, their high diversity and high abundance in *P. strigosa* through diel cycles suggest they could play a critical role in the holobiont. Other key members include *Ostreobium* spp., a photosynthetic endolithic symbiont that provides nutrient cycling and exchange of nitrogen and carbon, in a unique but often overlooked symbiosis ^14,107^.

The network analyses provide co-occurrence patterns as a proxy for microbial ecological interactions, which can be leveraged to hypothesize potential microbe-microbe interactions within the holobiont. The strongest association was observed between Symbiodiniaceae and the corallicolids, although the functional role of these apicomplexan coral dwellers remains unknown and are thus difficult to interpret. Co-occurrence networks also revealed diel structuring where one-third of the representative taxa showed 12-hour shifts, with microbes unique to day or night. Several bacterial members within these networks have been reported to perform roles in various key metabolic processes. For example, *Pseudomonas* and *Endozoicomonas* are capable of metabolizing dimethylsulfoniopropionate (DMSP), an organic sulfur compound produced by Symbiodiniaceae, using its cleavage products (methanethiol and acrylate) to support their growth ^106,120–124^. DMSP catabolism generates sulfur-containing antimicrobial compounds such as tropodithietic acid, a known *Vibrio* growth inhibitor ^95,125^, suggesting that these interactions may contribute to holobiont homeostasis. Daytime networks were enriched in taxa previously reported in carbohydrate metabolism, likely reflecting the influx of glucose and other photosynthates produced by Symbiodiniaceae during peak photosynthesis. Co-occurrences with *Defluviicoccus* (anaerobic glycogen-accumulating organisms; Bessarab et al. 2022), *Blautia*, *Bacteroides*, and *Parabacteroides*, all reported to catabolize carbohydrates into short-chain fatty acids such as propionate and acetate, suggest that these microbes are involved in processing algal photosynthates ^127,128^. At night, the presence of methylotrophic taxa (*Methylobacterium*, *Metholorubrum*) and methanogenic archaea implies the consumption of methanethiol, dimethyl sulfide (DMS), and glycolysis-derived byproducts ^129^. Taken together these co-occurrence patterns suggest that diel cycles influence microbial metabolic niches: enrichment of carbohydrate metabolizers during the day and methylotrophs and methanogens at night. This dynamic metabolic landscape likely sustains functional stability in the holobiont ^10^, capturing subtle but critical diel fluctuations in the coral ecosphere.

### Limitations of the study

Our study provides the first *in-situ* description of the transcriptomic and microbiome responses of *Pseudodiploria strigosa* in replicated natural diel cycles. These findings should be considered in the light of several limitations in our study design. Functional assignment is constrained by an annotation gap: most enrichment terms come from higher animal orthology, so their relevance to corals depends on how well they are conserved across lineages. Furthermore, we analyzed *de novo*-assembled transcripts rather than genome-modeled, curated genes; our totals likely include redundant isoforms, noise, and some contamination; and transcript-level regulation may not mirror gene- or protein-level regulation. In addition, gene expression patterns in eukaryotes are delayed due to a transcriptional lag of nascent *in-situ* mRNA and the rate of synthesis of mature mRNA ^130^. While the presence of a lag appears universal, its magnitude and duration vary, and these transcriptional delays must be considered when attributing function to time. Sampling coincided with a reef-wide spawning event involving several invertebrates, primarily from the second night into the third morning. *P. strigosa* typically spawns between July and September, with observations seen as late as October ^131^. Although we cannot determine whether reproduction-associated transcripts reflect *P. strigosa* preparing to spawn or having already spawned, we did not observe active spawning or visible gametes in the colony examined. Additionally, the *in-silico* metabarcoding approach relies on primers, which introduces bias common to all metabarcoding studies across different domains. For example, amplifying a short segment of the small subunit rRNA gene (using the 515F-806R primers) may overlook specific taxa due to insufficient nucleotide-level variation, and these primers are not universally effective for Archaea. Consequently, caution is warranted when interpreting microbial community structure, especially for the archaeal fraction. Extending these analyses across additional coral species will resolve holobiont dynamics at scale and clarify how homeostasis is maintained across divergent hosts.

## Acknowledgements

The authors would like to acknowledge and thank the CARMABI marine station in Willemstad, Curaçao. This project would not have been possible without the University of Miami’s start-up funding support and UM’s Frost Institute for Data Science and Computing resources towards the HPC cluster.

## Funding

University of Miami Start-up Funding (BAW, AMB, JdC)

Natural Sciences and Engineering Research Council of Canada Postgraduate Scholarship - Doctoral (BAW)

University of Miami IDSC Resources for Early Career Researchers (BAW)

## Author contributions

BAW and JdC conceptualized and designed the experiment. BAW and AMB conducted the sampling and processing of the samples. BAW and NK developed the bioinformatics pipeline and subsequent analyses. BAW wrote the manuscript, and JdC and ACB provided critical guidance and mentorship.

Conceptualization: BAW, JdC

Methodology: BAW, AMB, MJAV, JdC

Investigation: BAW, AMB, JdC

Visualization: BAW, NK, JdC

Supervision: ACB, JdC

Writing—original draft: BAW, JdC

Writing—review & editing: BAW, AMB, NK, MJAV, ACB, JdC

## Resource availability

### Lead contact

The lead contact for this study is Javier del Campo, who can be contacted at.

### Materials availability

The authors declare there are no novel reagents or materials generated from this study, and all resources are commercially available to reproduce this research.

### Data and code availability

All data is available through NCBI under BioProject XXXXXXX. All code used to analyze the data can be found at https://github.com/delCampoLab/diel_pstr.

## Competing Interests

Co-author Nicholas Kron is presently employed at Genevia Technologies, but all guidance and contributions were provided before employment. All other authors declare no competing interests.

## AI-Assisted Technologies

During the preparation of this work the authors used ChatGPT in the final stages for editing and proofreading. After using this tool/service, the authors reviewed and edited the content as needed and take full responsibility for the content of the publication.

## Supplemental Information

Supplementary Figure 1: (A) Map of Curaçao, (B) showing a close-up of the CARMABI research station and the specific sampling location marked in navy blue, labeled “Dreams”. (C) Colony of Pseudodiploria strigosa at the time of sample, and the image to the right is 6 months after sampling, illustrating the colony healed over where the samples were taken (lighter paling of the colony).

Supplementary Figure 2: Dot plots containing all (no filtering of “Human Diseases”) top enriched KEGG pathways grouped by time point (midnight 12 AM, dawn 6 AM, midday 12 PM, and dusk 6 PM) for *P. strigosa* and *Breviolum* sp. and the respective negative log_10_ adjusted p-value.

Supplementary Figure 3: Bar plots separated by time point (12 AM: “navy blue”; 6 AM: dawn, “grey”, 12 PM: midday, “yellow”; 6PM: dusk, “orange”) showing the rhythmic expression as (A) transcripts per million for *Pseudodiploria strigosa* and (B) raw counts for *Breviolum* sp. on the y-axis over time (h) on the x-axis. Within each bar plot, the left panel is through three days in six-hour intervals, and the right panel is the average over 24 hours. Above each plot is the transcript name, followed by EggNOG-derived description if present. On top of each bar plot is the associated GO term, if the transcript was annotated.

Supplementary Figure 4: Water samples collected adjacent to sampling coral colony, illustrating the prokaryotic (A-C, G) and microeukaryotic (D-F, H) communities. Alpha diversity metrics Shannon (A&D) and Richness (B&E), and beta diversity Aitchison distance PCA (C&F) with 95% confidence ellipses colored by the day the samples were collected. Community composition bubble plots showing phylogenetically sorted taxa at the genus level from the prokaryotic (Bacteria: G top panel, Archaea: G bottom panel) and microeukaryotic domains (Eukaryotes without Suessiales: H top panel, the proportion of microeukaryotic ASVs: H middle panel, and the Symbiodiniaceae: H bottom panel). Taxa are colored based on their class, with the exception of the Symbiodiniaceae, and the bubbles represent the relative abundance, where upper and lower bound standard deviation are reflected by bubble transparency. X-axis columns represent the mean relative abundance for each time point (midnight 12 AM, dawn 6 AM, midday 12 PM, and dusk 6 PM). All taxa shown are contributing >1% relative abundance across water samples, with the exception of the Archaea, which are >0.001%.

Supplementary Figure 5: (A-C) *Pseudodiploria strigosa* chloroplast-derived microeukaryotic alpha and beta diversity plots showing Shannon diversity (A), observed richness (B), and Aitchison distance PCA (C). Samples are grouped by time points: midnight (12 AM), dawn (6 AM), midday (12 PM), and dusk (6 PM). (D) Bubble plot showing phylogenetically sorted taxa for chloroplast-derived microeukaryotes at the genus level community composition on the x-axis by time of day on the y-axis. For context, the color of the bubble is indicated by the taxonomic subdivision. The size of the bubble indicates the average relative abundance by percent. In contrast, the lighter shade of the bubble represents the upper bound standard deviation, and the darkest represents the lower bound standard deviation. The “Below Threshold” category is the summation of the remaining taxa that do not contribute greater than 0.01% relative abundance. The “X”s beyond the taxonomy denote taxonomic placement from the highest taxonomic classification. (E) Complex heatmaps showing % read abundance of top 25 ASVs found in *P. strigosa* on the y-axis (Class; Family; Genus; ASV) by time on the x-axis (midnight: T12A, dawn: T6A, midday: T12P, and dusk: T6P), where samples are listed in order from day 1 (replicates 1 to 3) to day 3 (replicates 1 to 3).

Supplementary Figure 6: Complex heatmaps showing % read abundance of top 25 ASVs found in *Pseudodiploria strigosa* on the y-axis (Class; Family; Genus; ASV) by time on the x-axis (midnight: T12A, dawn: T6A, midday: T12P, and dusk: T6P), where samples are listed in order from day 1 (replicates 1 to 3) to day 3 (replicates 1 to 3). ASV taxonomy is listed for (A) prokaryome and (B) eukaryome, excluding Symbiodiniaceae reads.

Supplementary Figure 7: Differentially abundant (A) prokaryotic and (B) eukaryotic ASV-level taxa based on ANCOM-BC derived significance between midnight (12 AM) to midday (12 PM) and dawn (6 AM) to dusk (6 PM). Changes are based on significant log-fold change values using a false discover rate (FDR) adjusted p-value (alpha=0.05). Eukaryotic taxa used in analyses excluded Symbiodiniaceae to observe trends better.

## Notes

### Competing Interest Statement

The authors have declared no competing interest.

